# Ago2-seq identifies new microRNA targets for seizure control

**DOI:** 10.1101/777144

**Authors:** Morten T. Venø, Cristina R. Reschke, Gareth Morris, Niamh M. C. Connolly, Junyi Su, Yan Yan, Tobias Engel, Eva M. Jimenez-Mateos, Lea M. Harder, Dennis Pultz, Stefan J. Haunsberger, Ajay Pal, Braxton A. Norwood, Lara S. Costard, Valentin Neubert, Federico Del Gallo, Beatrice Salvetti, Vamshidhar R. Vangoor, Amaya Sanz Rodriguez, Juha Muilu, Paolo F. Fabene, R. Jeroen Pasterkamp, Jochen H.M. Prehn, Stephanie Schorge, Jens S. Andersen, Felix Rosenow, Sebastian Bauer, Jørgen Kjems, David C. Henshall

## Abstract

MicroRNAs (miRNAs) are short noncoding RNAs that shape the gene expression landscape, including during the pathogenesis of temporal lobe epilepsy (TLE). In order to provide a full catalog of the miRNA changes that happen during experimental TLE, we sequenced Argonaute 2-loaded miRNAs in the hippocampus of three different animal models at regular intervals between the time of the initial precipitating insult to the establishment of spontaneous recurrent seizures. The commonly upregulated miRNAs were selected for a functional *in vivo* screen using oligonucleotide inhibitors. This revealed anti-seizure phenotypes upon inhibition of miR-10a-5p, miR-21a-5p and miR-142a-5p as well as neuroprotection-only effects for inhibition of miR-27a-3p and miR-431-5p. Proteomic data and pathway analysis on predicted and validated targets of these miRNAs indicated a role for TGFβ signaling in a shared seizure-modifying mechanism. Together, these results identify functional miRNAs in the hippocampus and a pipeline of new targets for seizure control in epilepsy.

## Introduction

Temporal lobe epilepsy (TLE) is characterized by seizures arising from or involving the hippocampus and is the most common focal epilepsy syndrome in adults ^1^. TLE is frequently refractory to pharmacotherapy, often necessitating surgical resection of involved brain structures ^2^. The most common pathological finding within the removed hippocampus is select neuron loss and gliosis ^3^. Resected tissue from TLE patients also features neuroinflammation and remodeling of neuronal networks at both micro- and macroscopic scale ^4, 5^. Recent sequencing and array-based profiling of protein-coding transcripts and systems biology approaches have generated deep insights into the molecular pathophysiology and helped identify novel classes of molecule for therapeutic targeting ^6–9^.

MicroRNAs (miRNAs) are critical for shaping the gene expression landscape in the brain^10^. They are short noncoding RNAs that primarily function post-transcriptionally, conferring precision to cellular protein fluctuations ^11, 12^. Biogenesis of miRNAs involves nuclear processing of a primary transcript followed by terminal loop-processing in the cytoplasm, resulting in a miRNA duplex from which one strand is selected by an Argonaute (Ago) protein ^13^. Argonaute-2 (Ago2) is critically important for miRNA function, enriched in the hippocampus and, uniquely among Ago proteins, can directly cleave target RNAs ^14^. After miRNA loading and the formation of a RNA-induced silencing complex (RISC), potential mRNA targets are selected through imperfect base pairing between miRNA and mRNA ^15^. Upon identifying regions of sufficient complementarity, typically 7 – 8 nt matches between the miRNA and the 3’ untranslated region of the target mRNA, the RISC recruits further proteins, leading to translational repression or mRNA decay ^16^. Individual miRNAs often have multiple targets, increasing the scope for influencing several pathways or enhanced regulation of single pathways by multiple miRNAs, which may be an advantage for the treatment of TLE ^11, 12^.

Spatio-temporal changes to miRNA expression have been reported in the hippocampus following epileptogenic brain injuries and these persist in established epilepsy ^17, 18^. In parallel, *in vivo* deployment of oligonucleotide miRNA inhibitors (antagomirs) has demonstrated functional roles for a few miRNAs in seizure control and epileptogenesis ^19, 20^. It remains unknown how many more miRNAs may be suitable targets in epilepsy. Recent efforts have identified miRNAs dysregulated in TLE ^21–23^ but no study to date has focused on quantifying the amounts of functional Ago2-loaded miRNAs that are shared between TLE models. This is important since the specific enrichment for Ago2-loaded miRNAs provides greater coverage of the miRNA landscape and better predicts the regulatory potential of miRNAs ^24^.

Here, we performed small RNA sequencing of Ago2-loaded miRNAs from three different animal models across all phases of epilepsy development in two rodent species. Based on this resource, we deployed an *in vivo* antagomir screen and identified several novel anti-seizure and neuroprotective phenotypes among miRNAs that were up-regulated across the three models. Pathway analysis suggests TGFβ signaling as a potential overlapping mechanism shared between the target miRNAs. Our systems-level approach identifies an extensive class of miRNAs that may prove targetable for the treatment of seizures in human TLE.

## Results

### Ago2-seq provides a quantitative catalog of functionally-engaged miRNAs in experimental TLE

To identify shared functionally-engaged miRNAs at each phase of epilepsy development, we performed Ago2-immunoprecipitation followed by small RNA sequencing on hippocampal samples from three different rodent models of TLE in two species under continuous electroencephalogram (EEG) monitoring, sampling tissue at six time-points (intra-amygdala kainic acid (IAKA): N = 18 treated and 18 vehicle control; pilocarpine (PILO): N = 18 treated and 18 vehicle control; perforant path stimulation (PPS): N = 21 treated and 3 non-stimulated control, total N = 96; Figure 1A, Supplementary Data 1 and see *Methods*). This generated 1.44 billion small RNA reads of which up to 82% were miRNAs, with over 400 unique miRNAs detected per model (Figure 1B, Supplementary Data 2). There was exceptionally high concordance for levels among the most abundant miRNAs, with sequencing reads often differing by as little as 1 % across the three models (Figure 1C).

**Figure 1.**
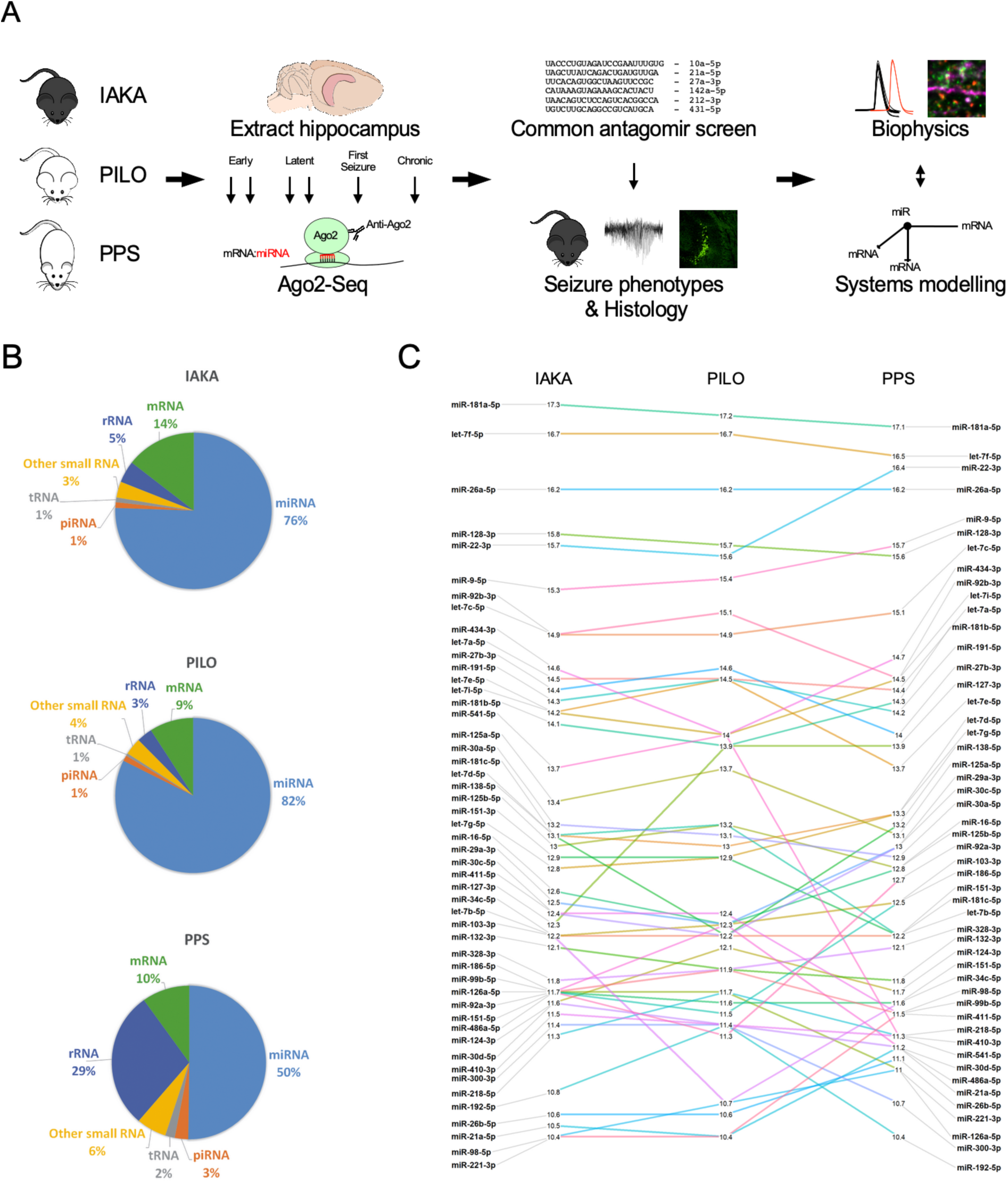
Experimental design and small RNA sequencing. *(**A**)* Schematic showing the full study design. 1) Three rodent models of epilepsy were generated: IAKA (intraamygdala kainic acid-induced status epilepticus in C57BL/6 mice), PILO (pilocarpine-induced status epilepticus in NMRI mice) and PPS (perforant pathway stimulation-induced hippocampal lesioning in Sprague-Dawley rats), 2) Hippocampi were extracted at six times-points and processed for Ago2 immunoprecipitation and small RNA sequencing (Ago2-seq). 3) Novel miRNAs with consistent up-regulation in all three models were selected for antagomir-based screen for anti-seizure phenotypes and neuroprotection. 4) Pathway modelling and biophysics approaches were used to investigate the function of the miRNAs. *(**B**)* The read mapping distribution for the three rodent models. Note, majority of small RNA reads mapped to miRNAs. *(**C**)* Expression of the top 50 miRNAs between the three models showing highly similar expression levels.

Induction of epilepsy led to significant changes in the abundance of approximately half of the detected miRNAs in the hippocampus in each model (Figure 2A and Supplementary Data 3). Expression changes showed disease stage-specific differences for individual miRNAs, including up- and down-regulation shortly after epileptogenic insult, on the day of first spontaneous seizure and chronic epilepsy, indicating that all phases of epilepsy development are associated with specific miRNA changes (Figure 2A, B, C). For miRNAs that originate from the same primary transcript, expression levels of miRNAs within the cluster often followed similar patterns (Figure 2C).

**Figure 2.**
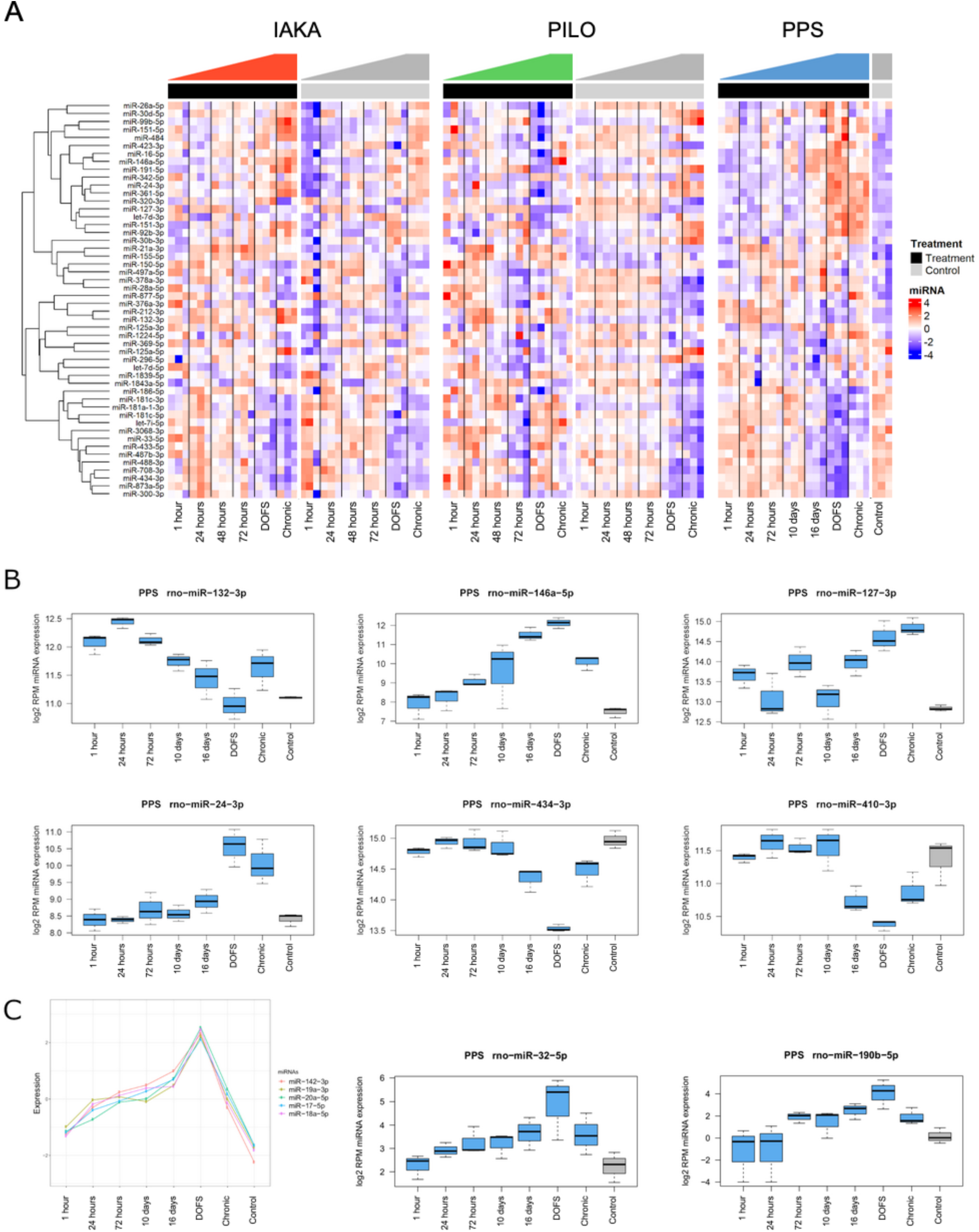
Extensive dysregulation of Ago2-loaded miRNAs across all phases of epilepsy development. *(**A**)* The 50 most significantly differentially expressed miRNAs are shown as a heatmap covering all samples from IAKA, PILO and PPS models. Top annotation shows epileptic animals as black and control animals as grey. Shown are z-scores of log2 transformed RPM values. *(**B**)* Examples of individual miRNA expression responses from the PPS model. Shown are miR-132 and miR-146 and potential novel epilepsy-associated miRNAs, miR-127, -24, - 434 and -410. *(**C**)* Clustering analysis shows that miRNAs from the miR-17∼92 cluster peak at DOFS. miR-142-3p also peaks at DOFS, though not transcribed from the miR-17∼92 cluster. Shown are also miR-32-5p and miR-190b-5p, both peaking at DOFS (day of first spontaneous seizure).

Next, we identified shared dysregulated (differential expression >25%) miRNAs across the three models, excluding low-abundance (<10 RPM) miRNAs for all time-points analysed (Figure 3A). Within the chronic epilepsy phase, the period most relevant to how a miRNA-based therapeutic might be used clinically (i.e. treating patients with pre-existing, refractory epilepsy)^17^, we found eight up- and one down-regulated miRNAs common to all three models (Figure 3B,C). This included miR-132-3p and miR-146a-5p, for which there is already significant functional data linking them to epilepsy ^25–28^, and six miRNAs (miR-10a-5p, miR-21a-3p, miR-27a-3p, miR-142a-5p, miR-212-3p and miR-431-5p) for which there is limited or no functional *in vivo* data linking to epilepsy (Figure 3C).

**Figure 3.**
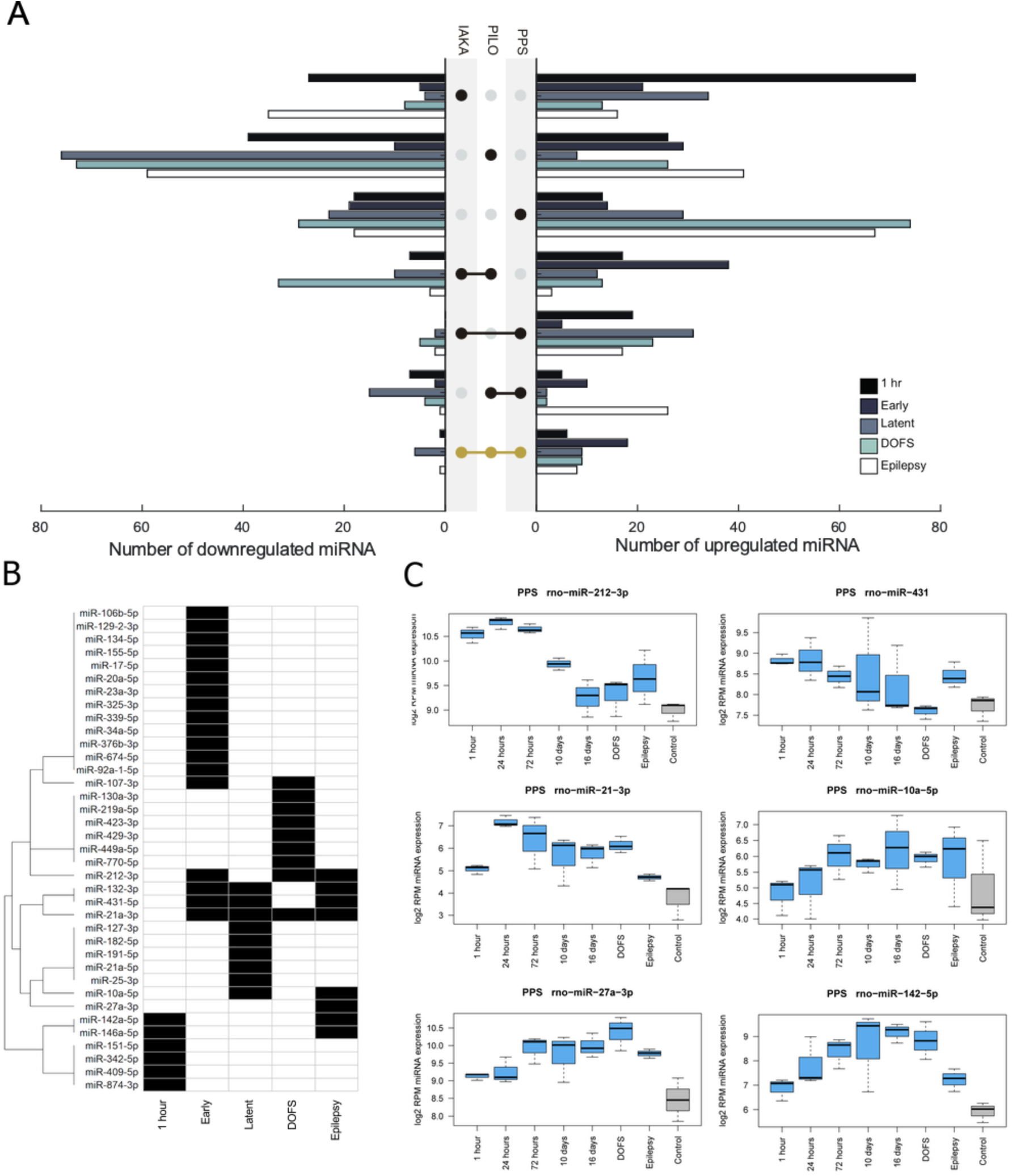
Identification of common-to-all model miRNAs. *(**A**)* Graphs show the overlap of up-and down-regulated miRNAs between the three models at various phases of epilepsy development. *(**B**)* The miRNAs with consistent up-regulation in all three models, common-to-all miRNAs, are further highlighted. *(**C**)* Examples of the expression data from the PPS model for the common-to-all miRNAs upregulated in chronic epilepsy (excluding miR-146a-5p and miR-132-3p).

### In vivo antagomir screening identifies three anti-seizure phenotypes

We hypothesized that the up-regulated miRNAs shared in the chronic epilepsy phase across the three models would be enriched for regulators of brain excitability. To test this, we assessed seizure responses after *in vivo* knock-down of miRNAs using locked nucleic acid (LNA)-modified oligonucleotide miRNA inhibitors (antagomirs). We excluded miR-132-3p and miR-146a-5p to prioritize miRNAs not previously linked to epilepsy, and excluded miR-21a-3p because it is not fully conserved in humans therefore limiting translational potential. Instead we selected the fully conserved miR-21a-5p, which also satisfied basal expression criteria and upregulation (at least 15%) in all three models. Mice received an intracerebroventricular injection of one of six targeting antagomirs, a scrambled antagomir or vehicle (PBS) 24 h before induction of status epilepticus by an intraamygdala microinjection of kainic acid (Figure 4A). This procedure ensures an optimal miRNA knockdown at the time of testing seizure responses ^29^. EEG recordings were used to assess seizure severity and brains were later processed to quantify irreversible hippocampal damage^29^.

**Figure 4.**
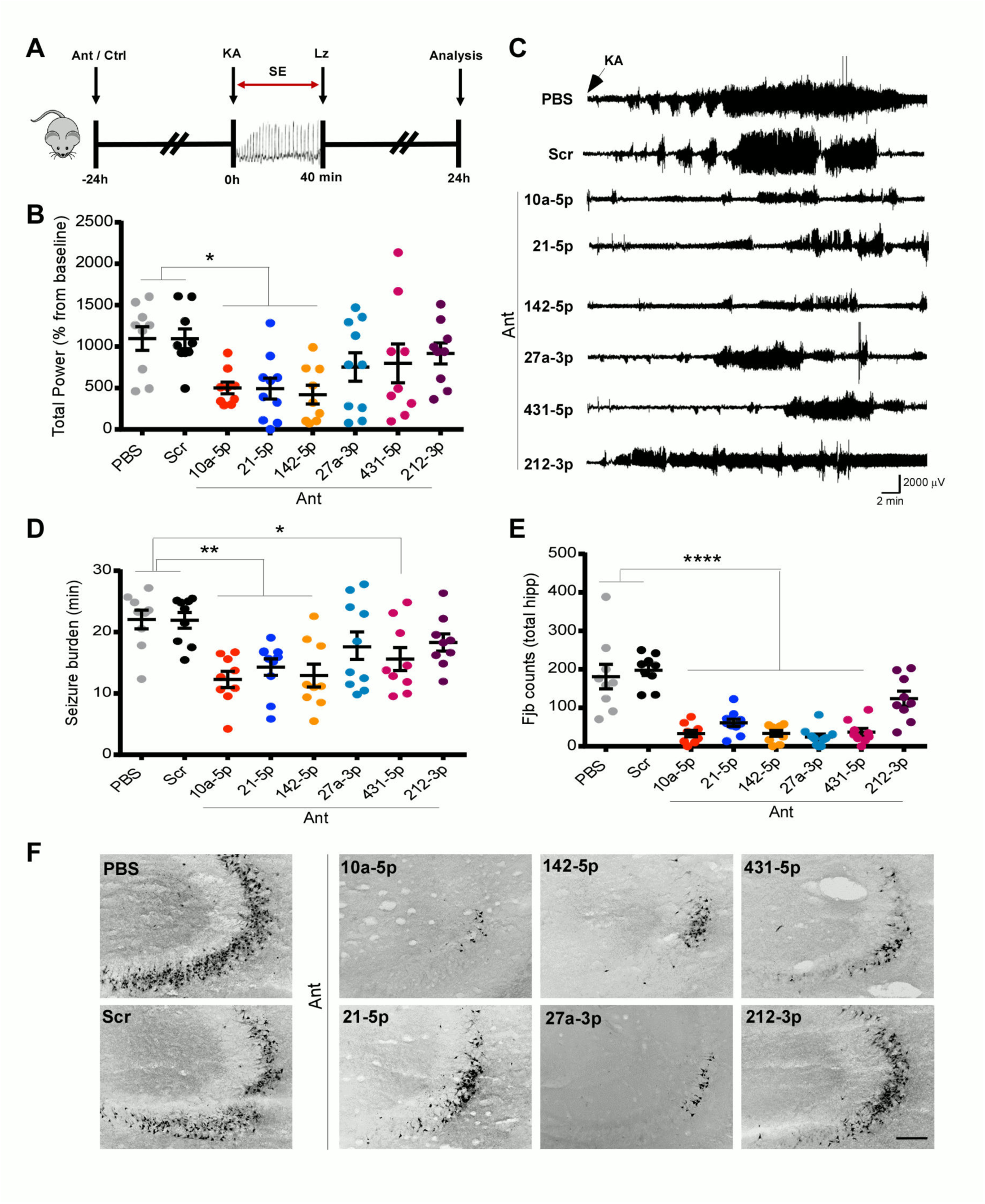
Seizure phenotype screening of antagomirs. ***(A)*** Schematic shows the experimental design. Briefly, mice were equipped for EEG recordings and underwent ICV injection of one of six antagomirs targeting miR-10a-5p, miR-21a-5p, miR-142a, miR-27a-3p, -5p, miR-431-5p, miR-212-3p and or controls (PBS or Scr). After 24 h, status epilepticus (SE) was induced by IAKA followed by lorazepam (Lz) to reduce mortality and morbidity. Hippocampal neuronal death was assessed at 24 h after SE. ***(B)*** Graph shows EEG total power during status epilepticus as a percentage of each animal’s own baseline data. Mice pre-treated with antagomirs for miR-10a-5p, miR-142a-5p, and miR-21a-5p displayed reduced seizure severity when compared to PBS or Scramble controls. ***(C)*** Representative traces show amplitude (µV) of EEG recordings over time (in min; starting from the IAKA injection) for each group. ***(D)*** Graph showing seizure burden (time in ictal activity) for each group. ***(E)*** Graph and ***(F)*** representative photomicrographs from the dorsal ipsilateral hippocampus of mice 24 h after status epilepticus, stained using the irreversible damage marker Fluoro-Jade B (FJB). Scale bars, 100 µm). All error bars shown as mean + S.E.M.. n = 9-10 / group; *P < 0.05, **P<0.01, ***P<0.001 compared either to PBS or Scr by One-Way ANOVA.

Seizure severity, as determined by analysis of EEG total power^29^, was significantly reduced during status epilepticus in mice pre-injected with antagomirs against miR-10a-5p, miR-21a-5p and miR-142a-5p (Figure 4B,C). Seizure burden, determined by measuring only ictal epileptiform activity^29^, was significantly reduced by the same antagomirs and was also significant for anti-miR-431-5p (Figure 4D). Analysis of the brains from mice killed 24 h after status epilepticus revealed significant neuroprotection for 5/6 of the antagomirs (those targeting miRNAs -10a-5p, -21a-5p, -27a-5p, -142a-5p and -431-5p), relative to controls (Figure 4E, F). These results suggest that a high proportion of the shared miRNAs upregulated in the chronic phase of experimental epilepsy may be mal-adaptive, contributing to enhanced network excitability and neuronal damage. Consequently, their targeting may offer novel approaches to seizure control.

### Knockdown of miR-10a-5p, -21a-5p and -142a-5p has limited biophysical and functional effects in naïve brains

Current anti-seizure drugs are associated with side effects including drowsiness that arise because of non-specific dampening of brain excitability ^1, 2^. To assess whether this was a risk for any of the antagomirs, we focused on the miRNAs which showed the most robust anti-seizure phenotypes when targeted (miR-10a-5p, miR-21a-5p and miR-142a-5p). We conducted a range of behavioral and electrophysiological assessments, originally developed for antagomir-injected rats,^30^ to report on possible adverse effects of miRNA inhibition *in vivo*. Performance of rats injected with each of the three antagomirs was normal in the novel object location test, although anti-miR-21a-5p caused a non-significant reduction in object discrimination (Figure 5A). We next prepared *ex vivo* brain slices from the same rats 2-4 days after antagomir injection, to coincide with maximal miRNA knockdown ^29^. Multiple electrophysiological measurements were normal in antagomir-injected animals including field response to Schaffer collateral stimulation (Figure 5B), paired pulse facilitation (Figure 5C) and pyramidal neuron biophysical properties (Figure 5D). We also stained hippocampal tissue sections for pre- and post-synaptic marker proteins. Immunofluorescent staining revealed that depletion of miR-142a-5p selectively reduced the size of the glutamatergic pre-synaptic marker VGLUT1, with no effect of the post-synaptic marker PSD-95. Anti-miR-10a-5p and anti-miR-21a-5p had no effect on either marker (Figure 5E). Taken together, these studies indicate that the anti-seizure antagomirs have no deleterious effects on hippocampal properties in naïve animals.

**Figure 5.**
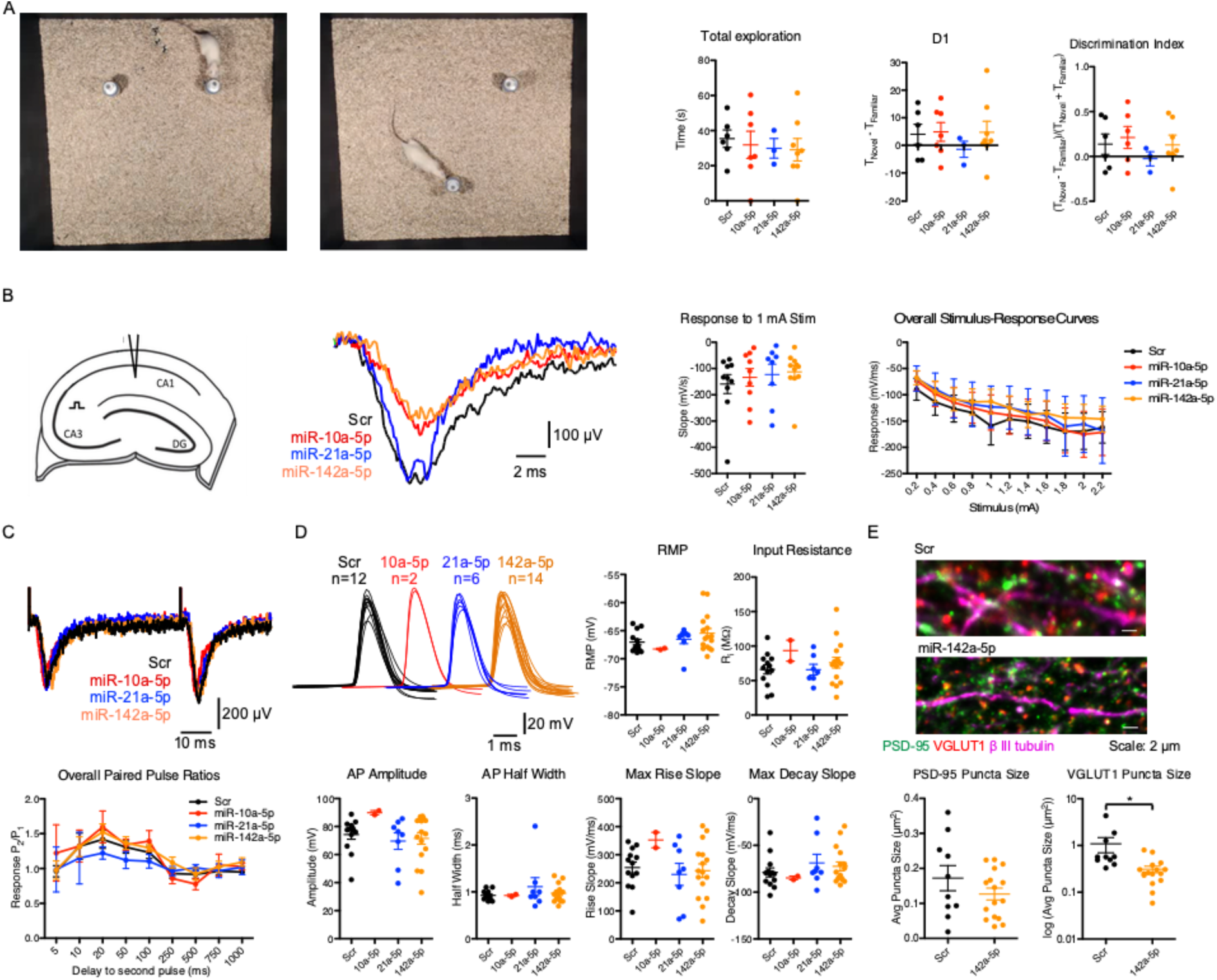
Antagomir effects on behavior and hippocampal biophysics in naïve animals. ***(A)*** Novel object location test - rats explored two identical objects for five minutes (*left image*) and were returned to the same arena after one hour with one object moved to a novel location (*right image*). Antagomirs did not cause a difference in total exploration time or any clear effect on absolute preference for the novel object (*D1 scatterplot*) or in discrimination index. ***(B)*** Stimulus-response curves - we stimulated the Schaffer collateral pathway in the hippocampus and recorded the population synaptic response in CA1 stratum radiatum (*left schematic*). Robust responses were observed in all treatment groups (*middle left panel*) and no significant differences in excitability were seen between groups (*right panels*). ***(C)*** Paired-pulse facilitation - we used the same electrode configuration as *(B)* but this time delivered two pulses (30% maximal response) at varying intervals. Robust facilitation was seen in all groups (*upper panel* - representative raw data for 50 ms stimulation interval) with no clear differences between groups (*lower panel*). ***(D)*** Single cell biophysics - we made current clamp recordings from CA1 pyramidal neurons and stimulated with a train of hyperpolarizing and depolarizing current steps. There were no differences in any passive properties (resting membrane potential (RMP) and input resistance scatterplots) or in properties of the threshold action potential (*upper left panel* - raw data for all recorded action potentials; lower scatterplots - properties of threshold action potentials). ***(E)*** Immunofluorescence for excitatory synaptic markers - staining for the excitatory pre-synaptic (VGLUT1) and postsynaptic (PSD-95) showed that ant-142 caused a selective reduction in VGLUT1 puncta size (*lower panel*: * - unpaired *t*-test on log-transformed data, p = 0.029).

### Convergence of targets and pathways regulated by the miRNAs

To investigate potential mechanisms of the seizure-modifying antagomirs, we developed a new miRNA-target interaction (MTI) database and focused on identifying convergent pathways for miR-10a-5p, miR-21a-5p and miR-142a-5p. The putative mRNA targets of the three miRNAs were identified using both predicted (miRDiP) ^31^ and experimentally validated (miRTarBase ^32^ TarBase ^33^) datasets. To reduce the risk of false-positives, we applied strict MTI filtering conditions based on miRDIP-assigned confidence levels and type of experimental validation (see Methods). The estimated MTIs for each miRNA, along with brain expression information for each putative target, are listed in Supplementary Data 4.

While each miRNA had many unique targets, the three seizure-related miRNAs (miR-10a-5p, miR-21a-5p and miR-142a-5p) shared 59 mRNA targets (Figure 6A, Table 1). Interestingly, SLC17a7 (VGLUT1) was predicted as a high confidence target of miR-142a-5p, in line with our experimental data (Figure 5F). 19 of the shared mRNA targets, including PTEN, were not targeted by miR-27a-3p or miR-431, indicating that these targets could be specific to the observed seizure-modifying effects (Figure 6A, Table 1). Moreover, 48 mRNAs targeted by >1 of the seizure-modifying miRNAs (of a total 525) have previously been associated with epilepsy, including GABA receptor, sodium, and potassium channel subunits (Table 2). *In situ* hybridization for miR-10a-5p, miR-21a-5p and miR-142a-5p suggested neuronal as well as glial expression, consistent with these targets (Supplementary Data 5).

**Figure 6:**
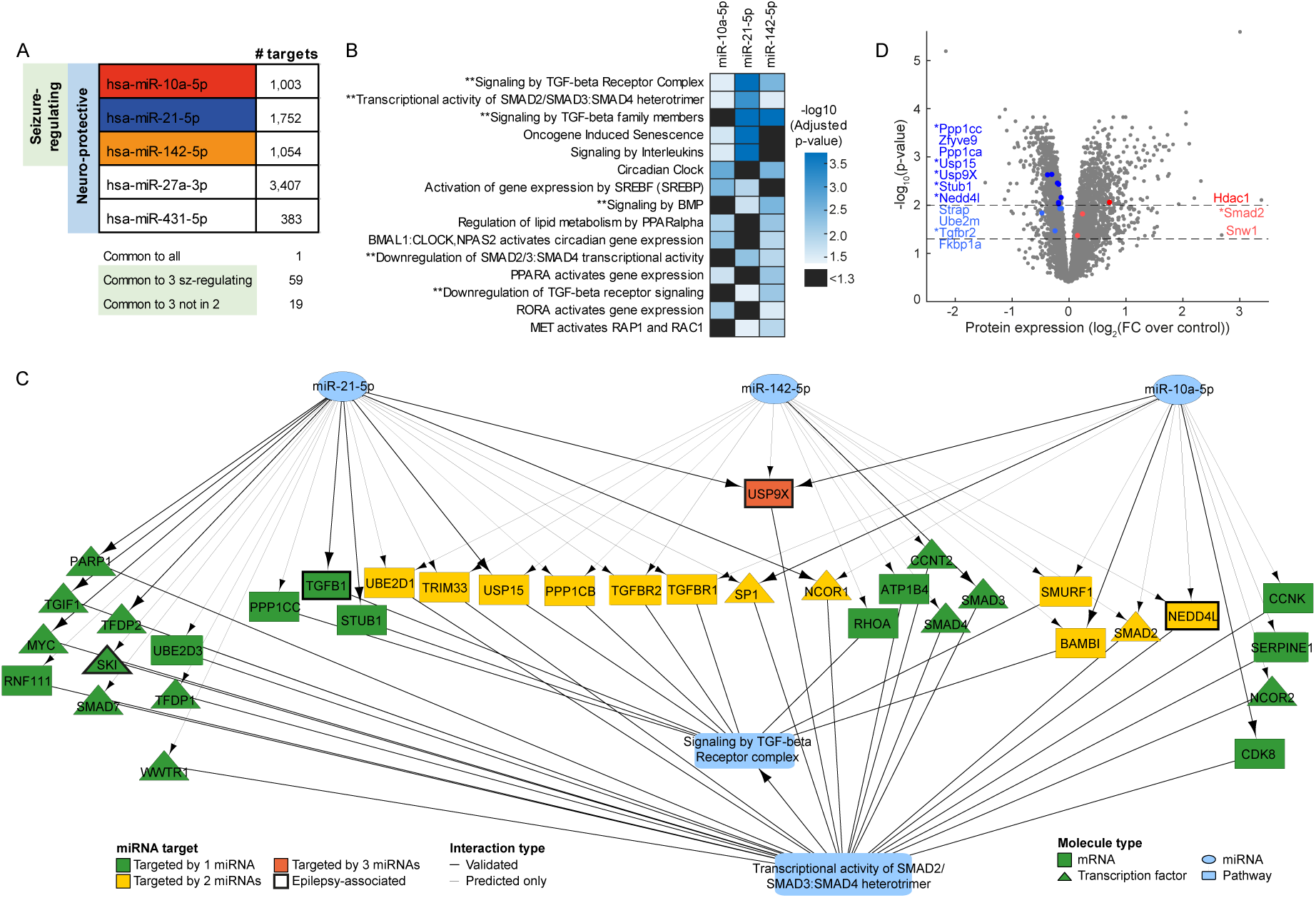
Target identification and pathway enrichment analysis identified TGFβ signaling as a potential convergent mechanism of the seizure-modifying miRNAs. (A) Number of mRNAs targeted by each miRNA. 1 mRNA (Thyroid Hormone Receptor Beta; THRB) is targeted by all 5 miRNA. 59 mRNA are targeted by the three seizure-modifying miRNAs, 19 of which are not targeted by miR-27a-3p nor miR-431 (see also Table 1). All targets are listed in Supplementary Data 5. (B) Significantly enriched Reactome pathways for each of the seizure-modifying miRNAs. ** indicates pathways associated with TGFβ signaling. (C) Wiring diagram depicting mRNA targets of the three seizure-modifying miRNAs that are involved in the reactome pathways ‘Signaling by TGF-beta receptor complex’ and ‘Transcriptional activity of SMAD2/SMAD3:SMAD4 heterotrimer’, illustrating the convergence of diverse miRNA targets at the pathway level. (D) Protein expression levels (normalized to control) of rat hippocampi isolated at the chronic time-point of the PPS model, (FC: foldchange). Proteins above the dashed lines (drawn at –log10(p-value) = 1.3, equivalent to p = 0.05 and at -log10(p-value) = 2, equivalent to p = 0.01) are considered significantly significant. Fold changes are shown on the x-axis with proteins involved in the TGF-β signaling pathways are highlighted in blue (downregulation) and red (upregulation). * denotes proteins which are targeted by miR-10a-5p, miR-21-5p and/or miR-142-5p, as depicted in panel C.

**Table 1.**
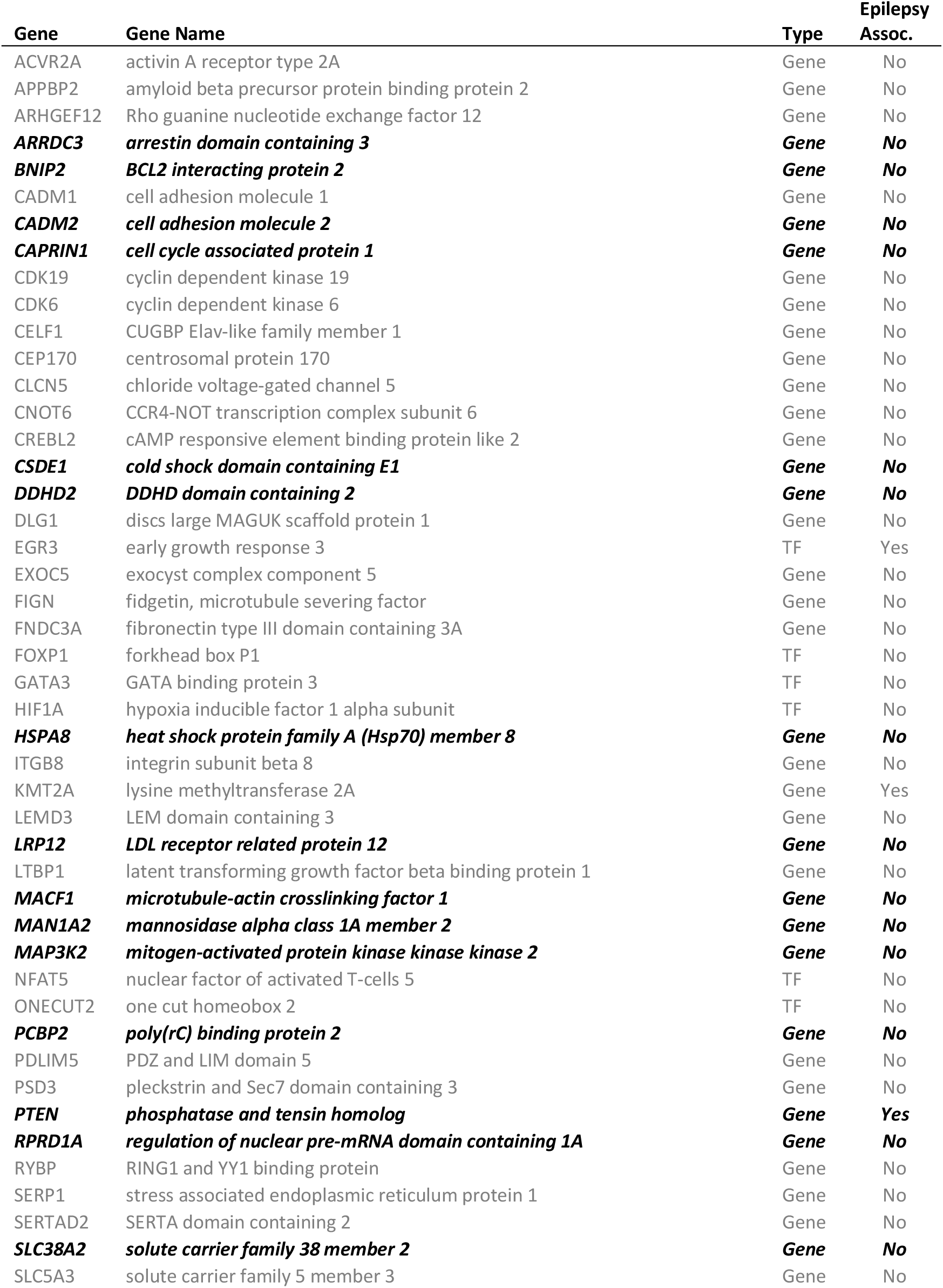
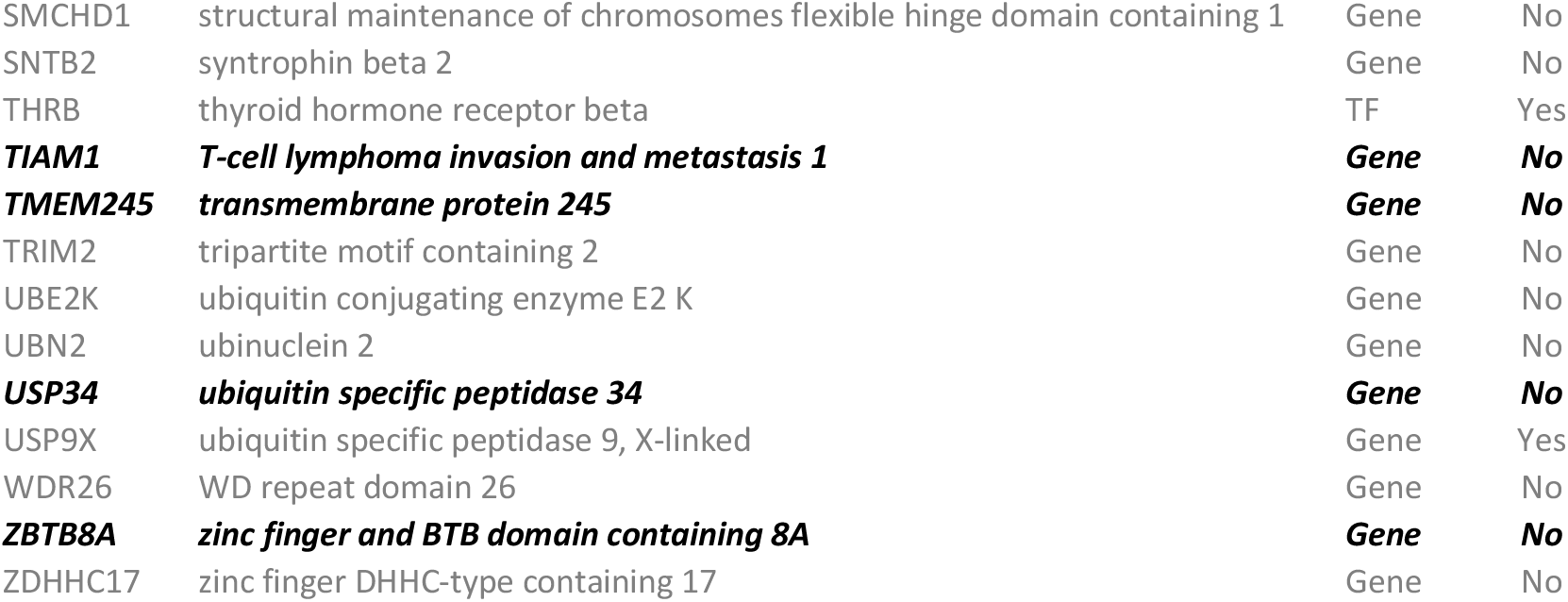
mRNAs targeted by all 3 seizure-modifying miRNAs (miR-142a-5p, miR-21a-5p, miR-10a-5p). mRNAs in ***black font*** are not targeted by miR-27a-3p nor miR-431, indicating that these targets may be specific to the observed seizure-modifying effects. See Supplementary Data 5 for more details. TF: transcription factor.

**Table 2.** mRNAs targeted by >1 seizure-modifying miRNA (miR-142a-5p, miR-21a-5p, miR-10a-5p) that have previously been associated with epilepsy. See Supplementary Data 5 for more details. TF: transcription factor

| gene | geneName | type | targeting miRNAs |
| --- | --- | --- | --- |
| ACTG1 | actin gamma 1 | Gene | 142-5p, 10-5p |
| BRWD3 | bromodomain and WD repeat domain containing 3 | Gene | 142-5p, 21-5p |
| CAMTA1 | calmodulin binding transcription activator 1 | Gene | 142-5p, 21-5p |
| CHL1 | cell adhesion molecule L1 like | Gene | 10-5p, 21-5p |
| DLG2 | discs large MAGUK scaffold protein 2 | Gene | 142-5p, 21-5p |
| DMD | dystrophin | Gene | 142-5p, 21-5p |
| DPYSL2 | dihydropyrimidinase like 2 | Gene | 10-5p, 21-5p |
| EGR3 | early growth response 3 | TF | 142-5p, 10-5p, 21-5p |
| FBXO28 | F-box protein 28 | Gene | 10-5p, 21-5p |
| FMR1 | fragile X mental retardation 1 | Gene | 142-5p, 21-5p |
| GABRB2 | gamma-aminobutyric acid type A receptor beta2 subunit | Gene | 10-5p, 21-5p |
| GATAD2B | GATA zinc finger domain containing 2B | Gene | 10-5p, 21-5p |
| HS2ST1 | heparan sulfate 2-O-sulfotransferase 1 | Gene | 142-5p, 21-5p |
| KCNA1 | potassium voltage-gated channel subfamily A member 1 | Gene | 10-5p, 21-5p |
| KCNJ10 | potassium voltage-gated channel subfamily J member 10 | Gene | 142-5p, 21-5p |
| KMT2A | lysine methyltransferase 2A | Gene | 142-5p, 10-5p, 21-5p |
| KRIT1 | KRIT1, ankyrin repeat containing | Gene | 142-5p, 21-5p |
| MBD5 | methyl-CpG binding domain protein 5 | Gene | 142-5p, 10-5p |
| MYT1L | myelin transcription factor 1 like | Gene | 142-5p, 10-5p |
| NEDD4L | neural precursor cell expressed, developmentally down-regulated 4-like, E3 ubiquitin protein ligase | Gene | 142-5p, 10-5p |
| NF1 | neurofibromin 1 | TF | 10-5p, 21-5p |
| NIN | ninein | Gene | 142-5p, 21-5p |
| NSD2 | nuclear receptor binding SET domain protein 2 | TF | 10-5p, 21-5p |
| PAFAH1B1 | platelet activating factor acetylhydrolase 1b regulatory subunit 1 | Gene | 10-5p, 21-5p |
| PLD1 | phospholipase D1 | Gene | 142-5p, 21-5p |
| PLP1 | proteolipid protein 1 | Gene | 10-5p, 21-5p |
| PTEN | phosphatase and tensin homolog | Gene | 142-5p, 10-5p, 21-5p |
| PTGS2 | prostaglandin-endoperoxide synthase 2 | Gene | 10-5p, 21-5p |
| PURA | purine rich element binding protein A | TF | 142-5p, 21-5p |
| RNF213 | ring finger protein 213 | Gene | 10-5p, 21-5p |
| SCARB2 | scavenger receptor class B member 2 | Gene | 10-5p, 21-5p |
| SCN3A | sodium voltage-gated channel alpha subunit 3 | Gene | 142-5p, 10-5p |
| SETD2 | SET domain containing 2 | Gene | 142-5p, 21-5p |
| SLC7A11 | solute carrier family 7 member 11 | Gene | 142-5p, 10-5p |
| SLC9A6 | solute carrier family 9 member A6 | Gene | 142-5p, 21-5p |
| SMC1A | structural maintenance of chromosomes 1A | Gene | 10-5p, 21-5p |
| SON | SON DNA binding protein | Gene | 142-5p, 10-5p |
| SOX5 | SRY-box 5 | TF | 142-5p, 21-5p |
| SPTAN1 | spectrin alpha, non-erythrocytic 1 | Gene | 10-5p, 21-5p |
| SYNJ1 | synaptojanin 1 | Gene | 142-5p, 10-5p |
| SYT14 | synaptotagmin 14 | Gene | 142-5p, 21-5p |
| TCF4 | transcription factor 4 | TF | 142-5p, 21-5p |
| THRB | thyroid hormone receptor beta | TF | 142-5p, 10-5p, 21-5p |
| UBR5 | ubiquitin protein ligase E3 component n-recognin 5 | Gene | 142-5p, 21-5p |
| USP9X | ubiquitin specific peptidase 9, X-linked | Gene | 142-5p, 10-5p, 21-5p |
| UTRN | utrophin | Gene | 142-5p, 21-5p |
| VDAC1 | voltage dependent anion channel 1 | Gene | 10-5p, 21-5p |
| ZMYND11 | zinc finger MYND-type containing 11 | TF | 10-5p, 21-5p |

We next performed Reactome pathway enrichment analysis on the predicted targets for each of the miRNAs, using targets expressed in the hippocampus, and found that 15 pathways were enriched for targets of >1 seizure-modifying miRNA (Figure 6B). Notably, six of these pathways are associated with TGFβ signaling, including the two pathways enriched in targets of all three miRNAs (‘R-HSA-170834: Signaling by TGF-beta Receptor Complex’ and its daughter pathway ‘R-HSA-2173793: Transcriptional activity of SMAD2/SMAD3:SMAD4 heterotrimer’). On further investigation, we noted that 35/73 genes involved in these two pathways are targeted by at least one of the seizure-modifying miRNAs (Figure 6C). Four of these mRNAs have been previously implicated in epilepsy, including Ubiquitin specific peptidase 9, X-linked (USP9X) ^34^, a gene targeted by all three miRNAs. To corroborate these systems-level predictions, we performed mass spectrometry proteomic analyses on rat hippocampi isolated at the chronic time-point of the PPS model. This identified significant changes in the expression of multiple proteins involved in TGFβ signaling. The main changes observed were down-regulation in the range of 0.7-0.9 FC (Figure 6D). This is consistent with the actions of miRNAs to fine tune protein levels of targets in the same pathway. Five of the seven significantly (p<0.01) downregulated proteins in the TGFβ signaling pathway, including Usp9x, are targeted by one or more of the three identified miRNAs as depicted in Figure 6C. Taken together, these results identify potential convergent miRNA target pathways underlying the anti-seizure effects of the miRNAs identified using functional screening across three *in vivo* animal models.

## Discussion

In the present study, we provide a comprehensive catalog of functional miRNA expression in the mouse and rat hippocampus and the changes that occur upon induction of epilepsy across three different models. Using this resource and an *in vivo* antagomir screening assay, we demonstrate that miRNAs that show consistent changes after spontaneous recurrent seizures in all three models are a rich source of targets for seizure modification. Intracerebral injection of anti-seizure antagomirs did not disrupt normal hippocampal functions. Finally, we provide evidence for pathways by which dysregulation of these miRNA may generate brain hyperexcitability. Together, these studies demonstrate how a systems-level approach can identify novel miRNA targets for the treatment of acute seizures or epilepsy.

By regulating the gene expression landscape and through their multi-targeting actions, miRNAs exert important effects on the excitable properties of neuronal networks underlying brain function ^10^. By extension, miRNAs represent potential targets for seizure control or disease-modification in epilepsy ^17^. The existence of conserved miRNA signatures in the development and maintenance of a seizure-prone state would provide important mechanistic insights and guide prioritization of miRNAs for therapeutic targeting. Here we undertook a coordinated effort, sequencing Ago2-bound miRNA to more accurately predict the regulatory potential of a given miRNA than by measuring overall miRNA levels in a sample^24^, covering three different models, two species and all stages from the initial precipitating insult to establishment of spontaneous recurrent seizures. The dataset contains robust statistics and fold change for individual miRNAs at each time point to illustrate expression variance and cross-model and cross-species comparisons. We found high concordance between the models and species in expression of known brain-enriched miRNAs, including miR-128-5p ^35^ and members of the let-7-family^36^ whereas no reads were detected for non-brain miRNAs such as miR-122-3p (liver-specific) and miR-208b-3p (heart-specific) ^37, 38^. The dataset features expected changes to neuronal activity-regulated miRNAs including miR-132-3p ^25^ and miRNAs that regulate cellular responses to tissue injury, such as apoptosis-associated miR-34a-5p ^39, 40^. Together, the results offer important advances over previous work which focused on predetermined miRNAs (e.g. microarray-based), lacked quantitative information on relative abundance and lacked functional relevance (non-Ago2-loaded miRNAs) ^21, 25, 41–45^. The Ago2-seq data provided in the current study are also an important companion to other databases on miRNA-epilepsy associations ^46^. The data complement, as well as reveal distinct profiles from, Ago2-seq analysis of neural precursors ^47^ and should interest researchers working on disease mechanisms for which there is shared pathophysiology, such as traumatic brain injury ^48^.

By employing a multi-model sequencing approach, we were able to demonstrate that there are shared miRNAs dysregulated at all phases in the development of epilepsy, up to and including the period of active chronic epilepsy. Most of the miRNA changes fell within a 1.5 – 3 fold range although some, including miR-142a-5p, displayed much larger fold changes. There was no apparent species or model-specific bias and numbers of shared miRNAs at the different stages of epilepsy development were quite similar, ranging from 6 - 18 among up-regulated miRNAs. We detected previously reported changes to miRNAs functionally linked to experimental epilepsy, including miR-22-3p ^49^, miR-129-5p ^50^, miR-134-5p ^25^, miR-146a-5p ^27^ and miR-324-5p ^51^. This indicates that Ago2-seq identifies robust miRNAs for targeting, a means to cross-compare between species and model, and a way to better prioritize miRNAs for functional assessment. A number of the miRNAs reported to be dysregulated in human TLE ^52–54^ were also differentially expressed in the chronic epilepsy state. This underscores the clinical relevance and translatability of our findings. It also invites additional predictions about human-dysregulated miRNAs which might be tested for function in animal models. The results extend evidence of a common miRNA signature in experimental epileptogenesis ^23^, contrasting conclusions from certain meta-analyses ^22^. Moreover, we report higher numbers of miRNAs and more differentially expressed miRNAs across these animal models than any previous epilepsy profiling study ^21, 25, 41–45^, indicating that miRNA dysregulation may impact on gene expression even more extensively than previously thought ^17^.

The potential for a miRNA-based therapeutic is gaining traction for disease modification in epilepsy ^5, 17^. LNA-based oligonucleotides as used here are particularly relevant for clinical translation as this backbone chemistry has been used in human trials of a miRNA-based therapy for hepatitis C ^55^. Here we show that robust anti-seizure and neuroprotective effects can be achieved by targeting multiple miRNAs commonly upregulated at the stage of chronic epilepsy. Notably, this included miRNAs for which there was no prior knowledge of a functional link to epilepsy. Our unbiased screen for anti-seizure phenotypes identified five antagomirs that protect the brain against prolonged seizures, of which those targeting miR-10a-5p, miR-21a-5p and miR-142a-5p had the most robust effects. This is a substantial addition to the number of miRNAs reported as potential targets for seizure control ^20^. It also suggests that many of the upregulated miRNAs in the chronic epilepsy phase may be suppressing targets that would otherwise oppose hyperexcitability. While the anti-seizure effects of targeting miR-10a-5p, miR-142a-5p and the neuro-protection associated with inhibition of miR-431-5p are novel, a recent study also found that targeting miR-21-5p could suppress seizures ^56^ Notably, our behavioral tests and biophysical analyses of the electrophysiological properties of hippocampus from antagomir-treated rodents showed no obvious impairments, indicating broad safety and suitability to enter preclinical development.

The regulatory potential of miRNAs is enhanced where there is convergence upon a small number of targets or pathways ^11, 12^. An important effort in the present study was to combine mRNA targets of all miRNAs (experimentally validated and predicted interactions) to build superior pathways, building in a high confidence threshold for target predictions. We found that mRNA targets of the three seizure-regulating miRNAs shared TGFβ and related SMAD signaling pathways as a potential overlapping seizure-modifying mechanism, while analyses at the protein level corroborated this pathway-level effect. This highlights that miRNA effects, while diverse at the level of individual miRNAs, can converge on common signaling pathways to exert complementary effects. TGFβ-signaling is known to be involved in epileptogenesis ^57^, and the seizure-suppressive effects of losartan, an AT1 receptor antagonist, are potentially mediated through TGFβ-signaling ^58^. Other overlapping targets included ion channels and PPARα-signaling pathways. Furthermore, several of the genes targeted by two or all three of the seizure-regulating miRNAs have previously been implicated in epilepsy. This includes USP9X ^34^ which is targeted by all three seizure-regulating miRNAs, was down-regulated in our proteomics analysis, and is also a component of the TGFβ-signaling pathway. However, the unique advantage of targeting miRNAs is the fact that by their nature of action, multiple genes within these signaling pathways are targeted.

There are some limitations and assumptions to consider in the present study. Some of the Ago2-bound miRNA pool may not be actively engaged with mRNA targets ^59^, Ago isoforms besides Ago2 may be important ^14^ and small RNA sequencing may over- or under-estimate the abundance of certain miRNA species ^60^. The use of already-epileptic animals, in which the target miRNA level would be increased, could have yielded larger effect sizes in our functional screen. While we restricted our functional studies to upregulated miRNAs, it is likely that seizure-regulating miRNAs are present among the downregulated miRNAs ^23^. Finally, adjustment of criteria for selecting miRNAs could yield additional miRNAs for functional studies. Indeed, several potentially new epilepsy-associated miRNAs not identified in multi-model or meta-analyses of miRNAs^21–23^ showed significant up- or down-regulation in two of the models here including highly-expressed miRNAs (thus likely to be functionally significant) such as miR-410-3p and miR-434-3p (down-regulated) and miR-24-3p and miR-127-3p (up-regulated) .

In summary, the present study generated a unique resource to explore the expression and dysregulation of miRNAs across multiple animal models of epilepsy and throughout the course of the disease. This systematic approach to discovery revealed a greater than previously anticipated, temporally-specific dysregulation of miRNAs in epilepsy and showed this to be a rich source of seizure-regulatory miRNA targets. We identified multiple additional miRNA targets for seizure control as well as identified potential mechanistic pathways. Together, these results reinforce and extend the evidence that miRNAs are a major class of regulatory element in epilepsy with therapeutic potential for seizure control.

## Methods

### Animal models of epilepsy

All animal experiments were performed in accordance with the European Communities Council Directive (2010/63/EU). All animals were housed in on-site barrier-controlled facilities having a 12 h light-dark cycle with ad libitum access to food and water.

Procedures in rats were approved by the local regulation authority (for Philipps University Marburg, Germany: Regierungspraesidium Giessen, 73/2013), or according to the Animals (Scientific Procedures) 1986 Act (UK). Male Sprague-Dawley rats (325–350 g; Charles River, Germany or 200-300 g; Harlan, UK) were used in all studies. Epilepsy was induced using the perforant pathway stimulation (PPS) model in rats, as described^61^. Animals received buprenorphine (0.2 mg/kg s.c.) and were anesthetized (isoflurane; 5% induction, 2-3% maintenance). Drill holes were prepared for electrode implantation and 3 fixing screws. An EEG transmitter (A3028E, Open Source Instruments, Inc., Watertown, MA, USA) was implanted into a skin pouch prepared at the left abdominal site of the rat. Stimulation electrodes (diameter 0.125 mm, Plastics One, Roanoke, VA, USA) were implanted bilaterally into the angular bundle of the PP (AP: immediately rostral of the lambdoid suture, ML: +/-4.5 mm lateral of the sagittal suture). Recording electrodes (diameter 0.25 mm, Plastics One, Roanoke, VA, USA) were implanted bilaterally into the DG (coordinates: 3.0 mm caudal from Bregma, +/-2.0 mm lateral from the sagittal suture). In order to determine the optimal dorso-ventral (DV) positioning of recording and stimulation electrodes, stimuli of 20 V at 0.5 Hz were applied with 0.5 Hz via the stimulation electrodes during implantation, and evoked potentials were recorded from the DG. Plastic connectors joined the electrodes with stimulation/recording equipment. After surgery, rats were allowed 1 week of recovery before PPS was initiated. The PPS protocol utilized a paradigm designed to evoke and maintain hippocampal seizure activity throughout the stimulation, but not convulsive status epilepticus^61^, which consisted of continuous, bilateral 2 Hz paired-pulse stimuli, with a 40 ms interpulse interval, plus a 10 second train of 20 Hz single-pulse stimuli delivered once per minute, generated by a S88 stimulator (Grass Instruments, West Warwick, USA). All pulses (0.1 ms duration) were delivered at 15-20 V. PPS was applied for 30 min on two consecutive days and for 8 h on the third day. As described previously, animals required only isoflurane (but no benzodiazepines or other injectable drugs) to terminate seizure activity which occurred occasionally immediately after the end of PPS^61^. Video and EEG were recorded continuously for up to 3 months. EEG recordings were performed with an Octal Data Receiver (A3027, Open Source Instruments, Inc., Watertown, MA, USA) with a sampling rate of 512 Hz. Data were recorded in NDF (Neuroscience Data Format) and converted to EDF (European Data Format) for visual analysis with EDFbrowser (version 1.57). Video recording was performed with infrared cameras (IC-7110W, Edimax Technology, Willich, Germany) and sampled with the SecuritySpy software (Ben Software Ltd., London, UK). The total EEG of all rats was screened visually for appearance of seizure patterns by experienced reviewers (LC, VN, BN, SB). In accordance with clinical practice in epileptology, seizure patterns were defined as rhythmic activity of at least 10 s which clearly broke background activity, contained epochs of high frequency spikes or spike-wave-complexes, and showed an evolution in frequency and amplitude. Video was used to clarify appearance of artefacts (e.g. chewing, scratching). Rats were killed under deep anesthesia (xylazine+ketamine) by transcardial perfusion with ice-cold 0.9 % NaCl solution at the following time points: 1 h, 24 h, 72 h, 10 d and 16 d after induction of epilepsy via PPS (epileptogenesis); within 1 d after the first spontaneous seizure (early epilepsy); 1 month after the first spontaneous seizure (chronic epilepsy). Control rats were killed on day 17 after surgery (corresponding to day 10 after PPS in the epilepsy group). Hippocampi were rapidly removed and snap frozen at -80 °C.

Procedures for inducing epilepsy using the intraamygdala kainic acid (IAKA) technique in mice were approved by the Research Ethics Committee of the Royal College of Surgeons in Ireland (REC-842), under license from the Health Products Regulatory Authority (AE19127/001), Dublin, Ireland. Adult male C57BL/6 mice (20 – 25 g, Harlan) were used, as described ^62^. Mice were anesthetized (isoflurane; 5% induction, 1–2% maintenance) and equipped for continuous EEG and video recordings using implantable EEG telemetry devices (Data Systems International). Transmitters (model F20-EET) which record bilateral EEG from the skull were implanted in a subcutaneous pocket at the time of cannula placement (on the dura mater following coordinates from bregma; IA: AP = -0.95 mm, L = +2.85 mm, V = 3.1 mm). The behavior of the animals was recorded using a video camera placed next to the cage. Continuous video-EEG data were acquired for each animal. After transmitter-cannula fitting, mice underwent intra-amygdala microinjection of kainic acid (IAKA; 0.3 µg in 0.2µl; Sigma-Aldrich, Ireland) to induce status epilepticus followed by intraperitoneal lorazepam (8 mg/kg) after 40 min to reduce morbidity and mortality. Mice were killed at 1 h, 24 h, 48 h, 72 h, the day of first spontaneous seizure (typically 3 – 5 d after status epilepticus) or at 2 weeks (chronic epilepsy). At the time of euthanasia, mice were deeply anesthetised with phenobarbital and transcardially perfused with ice-cold PBS to remove blood contaminants. Brains were rapidly removed and the entire hippocampus frozen and stored at -80°C.

Procedures for inducing epilepsy using the pilocarpine (PILO) model in mice were approved by the University of Verona research ethics committee under license from the Italian Ministry of Health (27/2014-PR). Adult male NMRI mice (Harlan) were fitted for DSI telemetry as above. After recovery, animals were given methylscopolamine (1 mg/kg) to block peripheral cholinergic actions and then after 30 min, given pilocarpine (300 mg/kg). Mice were and killed at 1 h, 24 h, 48 h, 72 h, the day of first spontaneous seizure (typically 1 -2 w after status epilepticus) or at 4 weeks (chronic epilepsy). Euthanasia and tissue preparation was as described above.

### Immunoprecipitation of Ago2, RNA extraction and sequencing (Ago2-seq)

Frozen hippocampi were allowed to thaw on ice. Thawed tissue was homogenised in 200 µl of IP buffer (300 mM NaCl, 5mM MgCl_2_, 0.1% NP-40, 50mM Tris-HCl pH 7.5, protease and RNase inhibitors) using plastic homogenising sticks until the tissue was completely homogenised. The homogenate was centrifuged at 16,000 g for 15 min at 4 °C to pellet nuclei and membranes. Supernatant (considered total cell lysate) was transferred to a clean Eppendorf tube. Bradford assay was performed to quantify protein content of total cell lysate. The lysate was pre-cleared by adding 10 µl of 50% Protein A/G beads (Santa Cruz Biotechnology, Germany) to 400 µg of protein lysate, final volume was adjusted to 1 ml using IP buffer and lysate was incubated rotating for 1 hour at 4 °C then centrifuged at 13,000 g for 5 min at 4°C to pellet the beads and supernatant was transferred to a new Eppendorf tube. 5 µg (5 µl of AGO-2 Cell Signalling Cat. #2897) antibody was added to pre-cleared cell lysate, vortexed and incubated rotating overnight at 4 °C. 20 µl of 50% A/G agarose beads were added to lysate-antibody solution and incubated rotating for 2 hours at 4 °C then centrifuged at 16,000 g for 15 min at 4°C and supernatant removed. The pellet was washed twice with 500 µl IP buffer by gently resuspending pellet, centrifuging at 16,000 g for 1 min at 4 °C and removing supernatant. Trizol RNA purification was performed after which pelleted RNA was dissolved in 12 µl dH20 and heated to 60 °C for 10 min. Purified RNA was stored at -80 °C until small RNA library preparation. 5 µl of purified RNA was prepared using TruSeq small RNA library preparation kit (Illumina), for rat samples using standard procedure and 12 PCR cycles, for mouse using half the amount of primers and reagents and 15 PCR cycles. Pippin prep (Sage Science) was used to size fractionate libraries to 140bp - 160bp size range. Library size and purity was validated on a Bioanalyzer 2100 (Agilent) using High Sensitivity DNA chip and the concentration was quantified using KAPA Library Quantification Kit. Prepared libraries were pooled as required and sequenced on a NextSeq500 (Illumina) at Exiqon.

### Analysis of small RNA sequencing data

FASTX-Toolkit was used to quality filter reads and cutadapt was used to remove adaptor sequences. Filtered reads were mapped using Bowtie to a list of datasets. First, reads were mapped to miRNAs from miRBase v21 allowing zero mismatches, but allowing for non-templated 3’ A and T bases. Reads not mapping to miRNAs were mapped against other relevant small RNA datasets: piRNA, tRNA, snRNA, snoRNA and Y RNA allowing one mismatch. The remaining unmapped reads were mapped to mRNA and rRNA datasets. MiRNAs were normalized as reads per million miRNA mapping reads (RPM). Statistical significance was calculated by One-Way ANOVA with FDR (Benjamini-Hochberg).

Common-to-all miRNAs were defined as having mean basal expression above 10 RPM, while exhibiting same-directional expression change of 25% or higher in all three models at key time points in each model. The time-points were: 1 h after induction of status epilepticus, 24 h (early epileptogenesis, in all models), latent period (72 h in IAKA and PILO models, 16 days in PPS model), day of first spontaneous seizure (DOFS; within the first 24 h of a first spontaneous seizure occurring), chronic epilepsy (2 weeks in the IAKA model, 4 weeks in the PILO model and 1 month in the PPS model). This approach allows for selection of miRNAs with similar expression trends in all three models at key functional periods. These common-to-all miRNAs are a strong vantage point for further validation.

### Systematic antagomir screening

Antagomir screening was performed in the IAKA mouse model according to previously described techniques ^63^. Mice were anesthetized and prepared with an additional guide cannula for intracerebroventricular antagomir injection (ICV: AP = +0.3 mm, L = +0.9 mm, V = 1.35 mm; from Bregma). Skull-mounted recording electrodes were placed and fixed with dental cement for EEG recordings. After recovery, mice were ICV injected with 0.5 nmol/2 µl of locked nucleic acid (LNA) oligonucleotide targeting: miR-10a-5p, miR-21a-5p, miR-27a-3p, miR-142a-5p, miR-212-3p or miR-431-5p. Control animals received a non-targeting scrambled version of the antagomir or PBS. Twenty-four hours later mice were connected to the lead socket of a swivel commutator, which was connected to an EEG (Grass TwiN digital EEG). A baseline recording was obtained followed by IAKA injection and continued for one hour. Mouse EEG data were analyzed and quantified using LabChart 8 software (ADInstruments, Oxford, U.K.) as described ^63^. Seizures were defined as high-amplitude (> 2 x baseline) high-frequency (> 5 Hz) polyspike discharges lasting > 5 seconds. EEG total power was plotted as percentage of baseline recording (each animal’s EEG power post seizure compared to its own baseline EEG) ^63^. Twenty-four hours after status epilepticus mice were transcardially perfused and brains sectioned for histopathological analysis of hippocampal damage. Seizure-induced neuronal damage was analyzed on 12 µm coronal sections at the level of medial hippocampus (AP = -1.70 mm) using Fluoro-Jade B (FJB) (Millipore Ireland B.V.) as described ^62^.

### In situ hybridization

Non-radioactive in situ hybridization (ISH) was performed as described previously ^64^. Except, hybridization was performed with 10 nM of double-DIG (3’ and 5’) - labelled locked nucleic acid (LNA) probes for miR-10a-5p, miR-142a-5p, miR-21a-5p and LNA-DIG Scramble probe (Exiqon) overnight at 50 °C followed by stringency washes at 55 °C. Four IAKA mice and two PBS mice with three sections per mouse were used.

### Behavioral testing and in vitro assay of effects of anti-miR-10a-5p, anti-miR-21a-5p and anti-miR-142a-5p

Stereotaxic injection for each miRNA knockdown was performed on adult male Sprague Dawley rats (weight range 270-380 g) as described previously ^30^. Briefly, rats were anaesthetized with isoflurane (5% induction, ∼2.5% maintenance) and given metacam (0.2 ml subcutaneous) prior to beginning surgery, followed by 0.15 ml buprenorphine and 2.5 ml saline (both subcutaneous) during recovery. We injected 2.5 nmol in 2 ml TE buffer of either anti-miR-10a-5p, anti-miR-21a-5p and anti-miR-142a-5p or Scr at the following co-ordinates (relative to bregma): AP -0.92 mm, L +1.3 mm, V 3.3 mm, to target the lateral ventricle. Treatments were blinded throughout all experimental procedures and analysis. Injection rate was controlled at 200 nL.min^-1^ and the needle was left in place for five minutes post-injection, to minimize backflow through the injection tract. Rats were allowed to recover from surgery with food and water freely available.

For the novel object location (NOL) test, rats were habituated to the behavioral arena (1m x 1m; Tracksys, Nottingham, UK) for five minutes each day over five days. On day six rats underwent stereotaxic surgery as described above. On day seven, rats completed the NOL test. Rats were allowed to explore two identical objects for five minutes. After one hour, rats were returned to the arena with one object moved to a novel location within the arena. Exploration was measured manually and defined as when the nose was within roughly 2 cm of the object, excluding time spent climbing on top of the object. Task performance was assessed using two measures: D1 (T_novel_-T_familiar_) and discrimination index ([T_novel_-T_familiar_]/ [Tnovel+Tfamiliar]).

*Ex vivo* brain slices were prepared between two and four days after surgery, to coincide with the maximal miRNA silencing effect ^30^. Rats were anaesthetized briefly with isoflurane and heavily with an i.p. injection of sodium pentobarbital, prior to transcardial perfusion with ice cold oxygenated sucrose ACSF slicing solution (in mM: 205 sucrose, 10 glucose, 26 NaHCO_3_, 1.2 NaH_2_PO_4_.H_2_O, 2.5 KCl, 5 MgCl_2_, 0.1 CaCl_2_). The brain was quickly extracted and sliced in 350 µm horizontal sections using a Campden 7000 smz slicer (Campden Instruments, Loughborough, UK). Slices for electrophysiology were held in a submerged style holding chamber filled with oxygenated recording ACSF (in mM: 125 NaCl, 10 glucose, 26 NaHCO_3_, 1.25 NaH_2_PO_4_.H_2_O, 3 KCl, 2 CaCl_2_, 1 MgCl_2_) and allowed to recover at room temperature for one hour before recording. Slices for immunohistochemistry were stored in 4% paraformaldehyde (PFA) at this point and processed as outlined below.

All slice electrophysiology was performed using a membrane chamber ^30^ perfused with oxygenated recording ACSF, heated to 34°C, at a rate of 16 mL.min^-1^. Electrophysiological data were recorded using an AxoClamp 700B amplifier (Molecular Devices, CA, USA), digitized at 10 kHz with a Power1401 (Cambridge Electronic Design, Cambridge, UK) and recorded using Signal software (Cambridge Electronic Design). For extracellular recordings, we stimulated the Schaffer Collateral pathway with a bipolar stimulating electrode and recorded the response in CA1 stratum radiatum using a glass microelectrode (∼5 MΩ) filled with recording ACSF. Patch clamp recordings used ∼5 MΩ glass microelectrodes filled with intracellular solution (in mM: 135 K-gluconate, 4 KCl, 10 HEPES, 4 Mg-ATP, 0.3 Na-GTP, 10 Na_2_-phosphocreatine; pH 7.3; 290 mOsm). All recordings were made with access resistance <20 MΩ and were rejected if action potentials did not overshoot 0 mV.

Slices for immunohistochemistry were fixed for 24 hrs in PFA before washing 3×5 mins in PBS. Slices were permeabilized for 2 hours at room temperature (RT) in PBS+0.5% triton and blocked for 1 hour at RT in PBS+3% BSA. Slices were incubated in primary antibodies overnight at 4 °C and washed again for 3×5 mins in PBS. Secondary antibodies were applied for 2 hours at RT before a final 3x5mins wash in PBS. Hoechst stain was added for 2 mins during the final wash. Slices were mounted using Fluoroshield (Sigma) and imaged using a Zeiss 710 confocal microscope (Carl Zeiss, Cambridge, UK).

### Bioinformatics – mRNA target identification, miRNA-target interaction (MTI) prioritization, and pathway enrichment analysis

We developed a novel Neo4j graph database that incorporates publicly available datasets of both predicted and experimentally validated miRNA-target interactions (MTIs). Predicted MTIs were downloaded from miRDiP V4.1, a database that integrates 30 prediction algorithms and calculates an MTI confidence score based on statistical inference [http://ophid.utoronto.ca/mirDIP/index.jsp#r]^31^. Experimentally validated MTIs were downloaded from miRTarBase V7 [http://mirtarbase.mbc.nctu.edu.tw/php/search.php#target]^32^ and TarBase V7.0 [http://carolina.imis.athena-innovation.gr/diana_tools/web/index.php?r=tarbasev8%2Findex]^33^. To ensure interoperability, all miRNA names were translated to miRBase V22 using miRNAmeConverter [http://www.systemsmedicineireland.ie/tools/mirna-name-converter/]^65^ and mRNA names were converted to official gene symbols using Ensembl V91. Non-human MTIs were excluded to constrain analyses to putative translatable mechanisms. Strict filtering criteria were devised to prioritize MTIs and reduce the risk of false-positive MTI identification - predicted MTIs were retained only if their miRDIP-assigned confidence levels were ‘Very High’, while experimentally validated MTIs were excluded if the only form of validation was high-throughput CLiP experiments performed prior to 2013, as these are considered less reliable due to poor antibody specificity and RNA contamination ^66^. Our database also incorporates baseline mRNA tissue expression information from the GTEx-EBI Expression Atlas [https://www.ebi.ac.uk/gxa/experiments/E-MTAB-5214/Results] and transcription factor information from TRRUST V2.0 [https://www.grnpedia.org/trrust/], ENCODE [http://amp.pharm.mssm.edu/Harmonizome/dataset/ENCODE+Transcription+Factor+Targets], and ChEA [http://amp.pharm.mssm.edu/Harmonizome/dataset/CHEA+Transcription+Factor+Targets]. mRNAs implicated in epilepsy were identified using an in-house database collating information from CARPEDB [http://carpedb.ua.edu/], epiGAD ^67^, Wang *et al* ^68^and curated epilepsy-genes from the Comparative Toxicogenomics Database (CTD) [http://ctdbase.org/ (Jan, 2019)]. ^69^ Pathway analysis was performed on Reactome pathways containing 10-500 genes by applying the cumulative hypergeometric distribution for p-value comparison ^70^. Pathways with corrected p-values < 0.05 (Benjamini-Hochberg) were considered significantly enriched.

### Proteomic analysis

Frozen rat hippocampi were resuspended in 200 µl lysis buffer (6M GdmCl, 10mM TCEP, 1× Complete protease inhibitor cocktail, 1× PhosSTOP inhibitor, 100 mM TEAB, benzonase) and dounced 30 times with a plastic douncer while kept on ice. Hereafter samples were boiled for 5 min and sonicated on ice using a probe sonicator with 10 sec on/ 10sec off for 2min total. Samples were then centrifuged at 13000 rpm for 3 min and the supernatant was added chloroacetamide to a final concentration of 20 mM followed by incubation in the dark for 30 min at room temperature (RT). Proteins where then digested adding first LysC and incubating for 30 min at RT and then diluting 10× using 100 mM TEAB followed by adding trypsin and incubating over night at 37°C with shaking. Both LysC and trypsin was added in a LysC/trypsin:protein ratio of 1:100.

Samples were centrifuged at 13000 rpm for 5 min and from the supernatant a volume corresponding to 50 µg peptide was transferred to a new eppendorff tube. Sets of 6 samples were labelled using the TMTsixplex™ isobaric label reagent. (ThermoFisher Scientific). The labeling reagent was prepared using the manufactures instructions to first equilibrate to RT, then dissolving the labeling reagent in 41 µl anhydrous acetonitrile for 5 min with occasional vortexing and finally centrifuging to gather the solution. The 41 µl labeling reagent was then added to each sample and incubated for 1 hour at RT. The reaction was quenched by incubating with 0.76 M lysine for 15 min. After quenching, a small amount was taken out from each sample, mixed and then tested by mass spectrometry analysis for labeling efficiency and mixing ratio. Hereafter the remaining samples were mixed in a 1:1 ratio.

The Stage-tips for the high-pH reverse phase fractionation was prepard by placing first two C18 disks in a p200 pipette tip, then adding a slurry of C18 beads (ReproSil-Pur, 3 µm, 120 Å) in methanol to create a layer of ∼1 cm of beads and adding one additional C18 disk in the top. The column was activated and equilibrated by washing one time in 150 µl buffer B (50% buffer A, 50% ACN) and then two times in 150 µl buffer A (20mM Ammonium hydroxide, pH 10). The labeled peptide sample was adjusted to pH 10 using buffer A and then loaded onto the activated and equilibrated stage-tip by spinning the sample through the column and collecting the flow-through. The column was washed 2 times with 150 µl buffer A and the washes were collected and combined with the flow-through. Next, peptides were eluted stepwise in 17 fractions by spinning 150 µl of buffer A containing increasing amounts of ACN for each step (3.5%, 4.5%, 6%, 8%, 10%, 10.5%, 11.5%, 13.5%, 15%, 16%, 17.5%, 18.5%, 19.5%, 21%, 27%, 50%, 80%) through the column. After collection, the liquid was evaporated completely in a Concentrator Plus (Eppendorf) and prepared for MS by resuspending in buffer A* (2% ACN, 0.1% TFA). Samples were analyzed using an Easy-nLC system coupled online to a Q Exactive HF mass spectrometer (Thermo Scientific) equipped with a nanoelectrospray ion source (Proxeon/Thermo Scientific). The peptides were eluted into the mass spectrometer during a chromatographic separation from a fused silica column packed in-house with 3 µm C18 beads (Reprosil, Dr. Maisch) using a 120 min gradient of buffer B (80% ACN, 0.5% acetic acid) with a flow rate of 250 nL/min. The Q Exactive HF was operated in positive ion mode with a top 12 data-dependent acquisition method where ions were fragmented by higher-energy collisional dissociation (HCD) using a normalized collisional energy (NCE)/ stepped NCE of 28 and 32. The resolution was set to 60,000 (at 400m/z), with a scan range of 300-1700 m/z and an AGC target of 3e6 for the MS survey. MS/MS was performed at a scan range of 100 – 2000 m/z using a resolution of 30,000 (at 400 m/z), an AGC target of 1e5, an intensity threshold of 1e5 and an isolation window of 1.2 m/z. Further parameters included an exclusion time of 45 sec and a maximum injection time for survey and MS/MS of 15 ms and 45 ms respectively.

Raw data from the LC-MS/MS analysis was processed using the MaxQuant software version 1.5.3.30 with default settings except from the following: In group specific parameters type was set to Reporter ion MS2 with 6plex TMT as isobaric labels. As variable modifications Oxidation (M), Acetyl (Protein N-term), and Deamidation (NQ) was added and in global parameters match between runs was selected. In MaxQuant, peak lists were searched against the rat UniProt database (including both swiss-prot and TrEMBL) released August 2016 using the build in search engine Andromeda.

Time profiles for each replicate were created by first normalizing to a common time point (day 16) present in each TMT experiment and then re-normalizing to the control. In this way each replicate time profile included protein ratios at each time point relative to the control. Differential expression was evaluated using a combined limma and rank product test to obtain q-values for each protein. Volcano plots were created by plotting log2 average ratios against – log10 q-values obtained in the above mentioned limma and rank product test.

### Statistics

For Ago2-seq statistical significance was calculated by one-way ANOVA with FDR (Benjamini-Hochberg). *In vivo* seizure screening used one-way ANOVA with Bonferroni post-hoc correction. *Ex vivo* slice experiments used unpaired *t*-test, one-way ANOVA, or mixed ANOVA, as appropriate. The statistical approach for pathway analysis and proteomics is detailed above.

### Data availability

Expression data have been submitted to the gene expression omnibus (GEO) under accession number GSE137473.

## Acknowledgements

We thank Lisa Anne Byrne for support with ethics and Anne Færch Nielsen, PhD for careful editing and suggestions on the manuscript. We would like to thank Nora Kalabrezi and Christian Siebert for help with EEG evaluation in the PPS model.

## Author contributions

MV, JS and YY performed Ago2 sequencing and statistical analyses. CRR performed *in vivo* antagomir studies and histological assessments in mice. GM and SS performed antagomir electrophysiology, behavior studies and immunofluorescent staining in rats. LMH, DP and JSA performed proteomics. SB, TE, E J-M, BS, FDG, BN, LC, VN generated brain samples. ASR performed EEG analyses. PFF, FR and DCH conceived and designed the animal studies, SJH, AP, NMCC, JHMP performed bioinformatics analysis, JK designed RNA sequencing studies. DCH wrote the initial manuscript and all authors contributed and approved the final manuscript.

---

**Supplementary Data 1.** Summary of EEG data for IAKA, PILO and PPS models.

**Animals sampled on day of first seizure**
| Animal ID | Species | Model | Latency to 1st Sz | No of Sz before hippocampus extracted | Time between final Sz and hippocampus extracted |
| --- | --- | --- | --- | --- | --- |
| RMK_DOFS_RH_85 | C57BL/6 mouse | IAKA | 73 hrs | 2 | 4 hrs |
| RMK_DOFS_RH_94 | C57BL/6 mouse | IAKA | 81 hrs | 1 | 4.5 hrs |
| RMK_DOFS_RH_95 | C57BL/6 mouse | IAKA | 73 hrs | 3 | 3.5 hrs |
| MRP07_RH_64 | Sprague Dawley rat | PPS | 33 days | 8 | 17 hrs |
| MRP07_RH_69 | Sprague Dawley rat | PPS | 12 days | 1 | 30.5 hrs |
| MRP07_RH_74 | Sprague Dawley rat | PPS | 19 days | 1 | - |
| VMP_DOFS_LH_1 | NMRI mouse | PILO | 86 hrs | 9 | 19 mins |
| VMP_DOFS_LH_2 | NMRI mouse | PILO | 47 hrs | 1 | 19 hrs |
| VMP_DOFS_LH_3 | NMRI mouse | PILO | 36 hrs | 1 | 10.5 hrs |

**Animals sampled in chronic epilepsy**
| Animal ID | Species | Model | Latency to 1st Sz | Total no of Sz | Total time in Sz | No of Sz in final 24 hrs | Time in Sz in last 24 hrs | Time between final Sz and hippocampus extracted |
| --- | --- | --- | --- | --- | --- | --- | --- | --- |
| RMK2W_RH_76 | C57BL/6 mouse | IAKA | 7 days | 25 | 502s | 0 | 0 | 72 hours |
| RMK2W_RH_101 | C57BL/6 mouse | IAKA | 3 days | 108 | 2014s | 11 | 229s | 4 hours |
| RMK2W_RH_106 | C57BL/6 mouse | IAKA | 3 days | 74 | 944s | 10 | 109.25s | 4 hours |
| MRP07_RH_37 | SD rat | PPS | 27 days | 4 | 161.7 | 0 | 0 | 101 hours |
| MRP07_RH_43 | SD rat | PPS | 11 days | 3 | 304 | 0 | 0 | 82 hours |
| MRP07_RH_75 | SD rat | PPS | 13 days | 9 | 318.4 | 0 | 0 | 135 hours |
| VMP4W_LH_1 | NMRI mouse | PILO | 88 hrs | - | - | - | - | - |
| VMP4W_LH_2 | NMRI mouse | PILO | 135 hrs | - | - | - | - | - |
| VMP4W_LH_3 | NMRI mouse | PILO | 17 days | - | - | - | - | - |

- : Data is unavailable

**Supplementary Data 2.** Summary of Ago2-seq reads.

**RNA species detected in Ago2 small RNA sequencing**
| RNA species | IAKA | PILO | PPS |
| --- | --- | --- | --- |
| miRNA | 75.5% | 82.4% | 50.2% |
| piRNA | 1.2% | 1.2% | 2.8% |
| tRNA | 1.0% | 0.5% | 2.0% |
| Other small RNA* | 3.3% | 3.6% | 6.3% |
| rRNA | 4.5% | 3.2% | 28.8% |
| mRNA | 14.4% | 9.1% | 9.9% |

\* Other small RNA refers to snoRNA, snRNA, yRNA, and small RNAs from Rfam (https://rfam.xfam.org/)

**Supplementary Data 3.** Expression of miRNAs in three rodent epilepsy models. This is too large to show here. A separate excel file has been generated.

**Supplementary Data 4.** MiRNA-Target Interactions for each miRNA, along with brain expression information for each target. This is too large to show here. A separate excel file has been generated.

**Supplementary Data 5.**
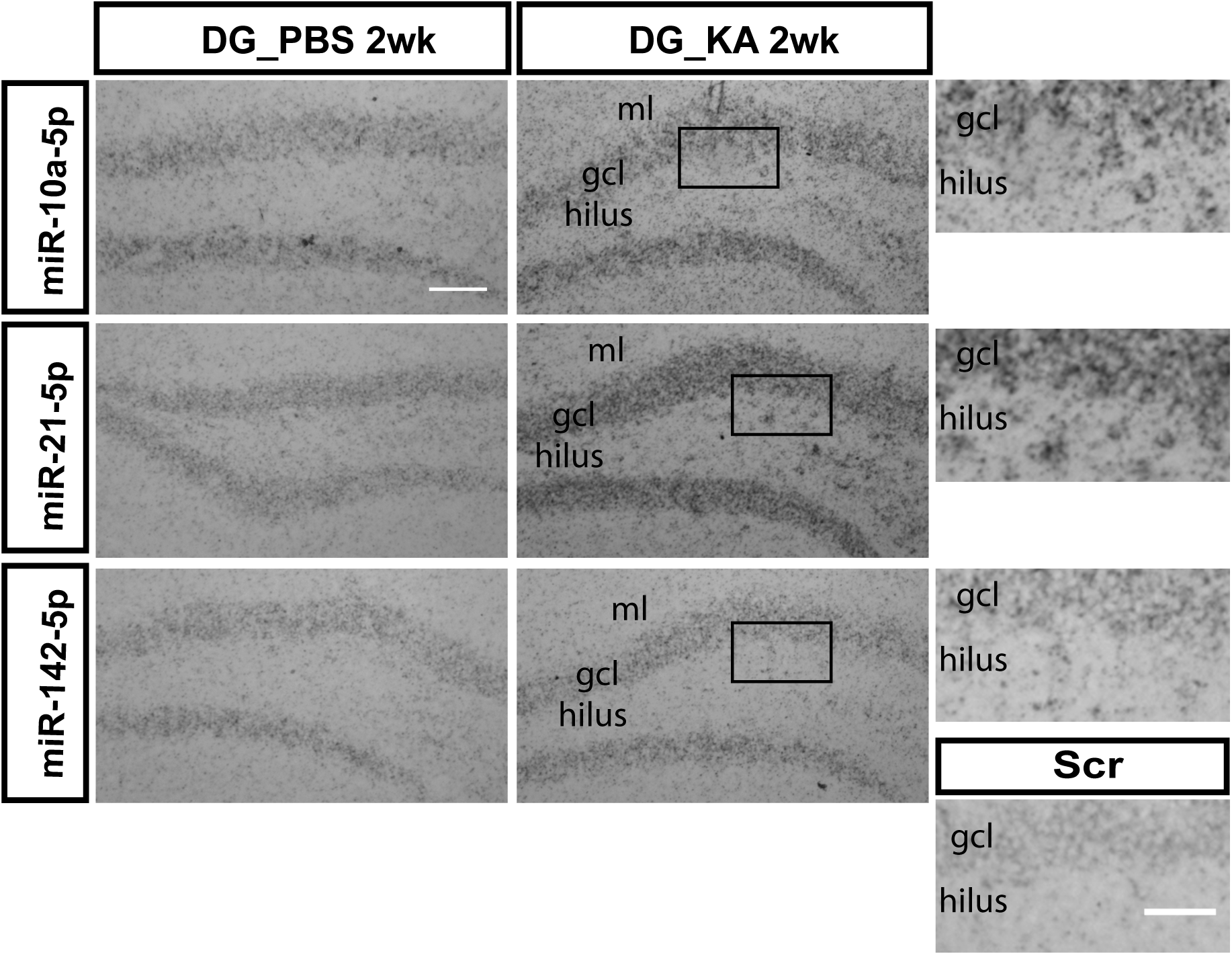
In situ hybridization for anti-seizure miRs. miRNA expression validation and cellular localization by *in situ* hybridization in IAKA mice at 2 weeks after injection. Low magnification images (main panels) show whole dentate gyrus (scale: 100 µm; scale for insets: 50 µm). All three miRs show stronger expression in KA injected mice compared to PBS injected mice.

## References

1. Schuele SU, Luders HO. Intractable epilepsy: management and therapeutic alternatives. Lancet Neurol 7, 514–524 (2008).

2. Kwan P, Schachter SC, Brodie MJ. Drug-resistant epilepsy. N Engl J Med 365, 919–926 (2011).

3. Blumcke I, et al. Histopathological findings in brain tissue obtained during epilepsy surgery. N Engl J Med 377, 1648–1656 (2017).

4. Pitkanen A, Lukasiuk K. Mechanisms of epileptogenesis and potential treatment targets. Lancet Neurol 10, 173–186 (2011).

5. Devinsky O, et al. Epilepsy. Nat Rev Dis Primers 4, 18024 (2018).

6. Gorter JA, et al. Potential new antiepileptogenic targets indicated by microarray analysis in a rat model for temporal lobe epilepsy. J Neurosci 26, 11083–11110 (2006).

7. McClelland S, et al. The transcription factor NRSF contributes to epileptogenesis by selective repression of a subset of target genes. Elife 3, e01267 (2014).

8. Johnson MR, et al. Systems genetics identifies Sestrin 3 as a regulator of a proconvulsant gene network in human epileptic hippocampus. Nat Commun 6, 6031 (2015).

9. Srivastava PK, et al. A systems-level framework for drug discovery identifies Csf1R as an anti-epileptic drug target. Nat Commun 9, 3561 (2018).

10. Kosik KS. The neuronal microRNA system. Nat Rev Neurosci 7, 911–920 (2006).

11. Schmiedel JM, et al. Gene expression. MicroRNA control of protein expression noise. Science 348, 128–132 (2015).

12. Bartel DP. Metazoan MicroRNAs. Cell 173, 20–51 (2018).

13. Ha M, Kim VN. Regulation of microRNA biogenesis. Nat Rev Mol Cell Biol 15, 509–524 (2014).

14. Czech B, Hannon GJ. Small RNA sorting: matchmaking for Argonautes. Nat Rev Genet 12, 19–31 (2011).

15. Chandradoss SD, Schirle NT, Szczepaniak M, MacRae IJ, Joo C. A dynamic search process underlies microRNA targeting. Cell 162, 96–107 (2015).

16. Gebert LFR, MacRae IJ. Regulation of microRNA function in animals. Nat Rev Mol Cell Biol, (2018).

17. Henshall DC, et al. MicroRNAs in epilepsy: pathophysiology and clinical utility. Lancet Neurol 15, 1368–1376 (2016).

18. Brennan GP, Henshall DC. microRNAs in the pathophysiology of epilepsy. Neurosci Lett 667, 47–52 (2018).

19. van Rooij E, Kauppinen S. Development of microRNA therapeutics is coming of age. EMBO Mol Med 6, 851–864 (2014).

20. Henshall DC. Manipulating microRNAs in murine models: Targeting the multi-targeting in epilepsy. Epilepsy Curr 17, 43–47 (2017).

21. Kretschmann A, et al. Different microRNA profiles in chronic epilepsy versus acute seizure mouse models. J Mol Neurosci 55, 466–479 (2015).

22. Srivastava PK, et al. Meta-analysis of microRNAs dysregulated in the hippocampal dentate gyrus of animal models of epilepsy. eNeuro 4, (2017).

23. Cava C, Manna I, Gambardella A, Bertoli G, Castiglioni I. Potential Role of miRNAs as Theranostic Biomarkers of Epilepsy. Mol Ther Nucleic Acids 13, 275–290 (2018).

24. Flores O, Kennedy EM, Skalsky RL, Cullen BR. Differential RISC association of endogenous human microRNAs predicts their inhibitory potential. Nucleic Acids Res 42, 4629–4639 (2014).

25. Jimenez-Mateos EM, et al. miRNA Expression profile after status epilepticus and hippocampal neuroprotection by targeting miR-132. Am J Pathol 179, 2519–2532 (2011).

26. Huang Y, Guo J, Wang Q, Chen Y. MicroRNA-132 silencing decreases the spontaneous recurrent seizures. Int J Clin Exp Med 7, 1639–1649 (2014).

27. Iori V, et al. Blockade of the IL-1R1/TLR4 pathway mediates disease-modification therapeutic effects in a model of acquired epilepsy. Neurobiol Dis 99, 12–23 (2017).

28. Tao H, et al. Intranasal Delivery of miR-146a Mimics Delayed Seizure Onset in the Lithium-Pilocarpine Mouse Model. Mediators Inflamm 2017, 6512620 (2017).

29. Jimenez-Mateos EM, et al. Silencing microRNA-134 produces neuroprotective and prolonged seizure-suppressive effects. Nat Med 18, 1087–1094 (2012).

30. Morris G, Brennan GP, Reschke CR, Henshall DC, Schorge S. Spared CA1 pyramidal neuron function and hippocampal performance following antisense knockdown of microRNA-134. Epilepsia in press, (2018).

31. Tokar T, et al. mirDIP 4.1-integrative database of human microRNA target predictions. Nucleic Acids Res 46, D360–D370 (2018).

32. Chou CH, et al. miRTarBase update 2018: a resource for experimentally validated microRNA-target interactions. Nucleic Acids Res 46, D296–D302 (2018).

33. Karagkouni D, et al. DIANA-TarBase v8: a decade-long collection of experimentally supported miRNA-gene interactions. Nucleic Acids Res 46, D239–D245 (2018).

34. Paemka L, et al. Seizures are regulated by ubiquitin-specific peptidase 9 X-linked (USP9X), a de-ubiquitinase. PLoS Genet 11, e1005022 (2015).

35. Tan CL, et al. MicroRNA-128 governs neuronal excitability and motor behavior in mice. Science 342, 1254–1258 (2013).

36. Shinohara Y, et al. miRNA profiling of bilateral rat hippocampal CA3 by deep sequencing. Biochem Biophys Res Commun 409, 293–298 (2011).

37. Sempere LF, Freemantle S, Pitha-Rowe I, Moss E, Dmitrovsky E, Ambros V. Expression profiling of mammalian microRNAs uncovers a subset of brain-expressed microRNAs with possible roles in murine and human neuronal differentiation. Genome Biol 5, R13 (2004).

38. Ludwig N, et al. Distribution of miRNA expression across human tissues. Nucleic Acids Res 44, 3865–3877 (2016).

39. Chang TC, et al. Transactivation of miR-34a by p53 broadly influences gene expression and promotes apoptosis. Mol Cell 26, 745–752 (2007).

40. Sano T, Reynolds JP, Jimenez-Mateos EM, Matsushima S, Taki W, Henshall DC. MicroRNA-34a upregulation during seizure-induced neuronal death. Cell Death Dis 3, e287 (2012).

41. Hu K, et al. Expression profile of microRNAs in rat hippocampus following lithium-pilocarpine-induced status epilepticus. Neurosci Lett 488, 252–257 (2011).

42. Bot AM, Debski KJ, Lukasiuk K. Alterations in miRNA levels in the dentate gyrus in epileptic rats. PLoS One 8, e76051 (2013).

43. Gorter JA, et al. Hippocampal subregion-specific microRNA expression during epileptogenesis in experimental temporal lobe epilepsy. Neurobiol Dis 62, 508–520 (2014).

44. Li MM, et al. Genome-wide microRNA expression profiles in hippocampus of rats with chronic temporal lobe epilepsy. Sci Rep 4, 4734 (2014).

45. Roncon P, et al. MicroRNA profiles in hippocampal granule cells and plasma of rats with pilocarpine-induced epilepsy - comparison with human epileptic samples. Sci Rep 5, 14143 (2015).

46. Mooney C, Becker BA, Raoof R, Henshall DC. EpimiRBase: a comprehensive database of microRNA-epilepsy associations. Bioinformatics 32, 1436–1438 (2016).

47. Liu XS, et al. Identification of miRNomes associated with adult neurogenesis after stroke using Argonaute 2-based RNA sequencing. RNA Biol 14, 488–499 (2017).

48. Liou AK, Clark RS, Henshall DC, Yin XM, Chen J. To die or not to die for neurons in ischemia, traumatic brain injury and epilepsy: a review on the stress-activated signaling pathways and apoptotic pathways. Prog Neurobiol 69, 103–142 (2003).

49. Jimenez-Mateos EM, et al. MicroRNA targeting of the P2X7 purinoceptor opposes a contralateral epileptogenic focus in the hippocampus. Scientific Reports 5, e17486 (2015).

50. Rajman M, et al. A microRNA-129-5p/Rbfox crosstalk coordinates homeostatic downscaling of excitatory synapses. EMBO J, (2017).

51. Gross C, et al. MicroRNA-mediated downregulation of the potassium channel Kv4.2 contributes to seizure onset. Cell Rep 17, 37–45 (2016).

52. Bencurova P, et al. MicroRNA and mesial temporal lobe epilepsy with hippocampal sclerosis: Whole miRNome profiling of human hippocampus. Epilepsia 58, 1782–1793 (2017).

53. Miller-Delaney SF, et al. Differential DNA methylation profiles of coding and non-coding genes define hippocampal sclerosis in human temporal lobe epilepsy. Brain 138, 616–631 (2015).

54. Kan AA, et al. Genome-wide microRNA profiling of human temporal lobe epilepsy identifies modulators of the immune response. Cell Mol Life Sci 69, 3127–3145 (2012).

55. Janssen HL, et al. Treatment of HCV infection by targeting microRNA. N Engl J Med 368, 1685–1694 (2013).

56. Tang C, et al. Targeting of microRNA-21-5p protects against seizure damage in a kainic acid-induced status epilepticus model via PTEN-mTOR. Epilepsy Res 144, 34–42 (2018).

57. Cacheaux LP, et al. Transcriptome profiling reveals TGF-beta signaling involvement in epileptogenesis. J Neurosci 29, 8927–8935 (2009).

58. Bar-Klein G, et al. Losartan prevents acquired epilepsy via TGF-beta signaling suppression. Ann Neurol 75, 864–875 (2014).

59. La Rocca G, et al. In vivo, Argonaute-bound microRNAs exist predominantly in a reservoir of low molecular weight complexes not associated with mRNA. Proc Natl Acad Sci U S A 112, 767–772 (2015).

60. Leshkowitz D, Horn-Saban S, Parmet Y, Feldmesser E. Differences in microRNA detection levels are technology and sequence dependent. RNA 19, 527–538 (2013).

61. Norwood BA, et al. Classic hippocampal sclerosis and hippocampal-onset epilepsy produced by a single “cryptic” episode of focal hippocampal excitation in awake rats. J Comp Neurol 518, 3381–3407 (2010).

62. Mouri G, et al. Unilateral hippocampal CA3-predominant damage and short latency epileptogenesis after intra-amygdala microinjection of kainic acid in mice. Brain Res 1213, 140–151 (2008).

63. Reschke CR, et al. Potent anti-seizure effects of locked nucleic acid antagomirs targeting miR-134 in multiple mouse and rat models of epilepsy. Mol Thera 6, 45–56 (2017).

64. Vangoor VR, et al. Antagonizing increased miR-135a levels at the chronic stage of experimental TLE reduces spontaneous recurrent seizures. J Neurosci, (2019).

65. Haunsberger SJ, Connolly NM, Prehn JH. miRNAmeConverter: an R/Bioconductor package for translating mature miRNA names to different miRBase versions. Bioinformatics, (2016).

66. Moore MJ, Zhang C, Gantman EC, Mele A, Darnell JC, Darnell RB. Mapping Argonaute and conventional RNA-binding protein interactions with RNA at single-nucleotide resolution using HITS-CLIP and CIMS analysis. Nat Protoc 9, 263–293 (2014).

67. Tan NC, Berkovic SF. The Epilepsy Genetic Association Database (epiGAD): analysis of 165 genetic association studies, 1996-2008. Epilepsia 51, 686–689 (2010).

68. Wang J, et al. Epilepsy-associated genes. Seizure 44, 11–20 (2017).

69. Davis AP, et al. The Comparative Toxicogenomics Database: update 2019. Nucleic Acids Res 47, D948–D954 (2019).

70. Boyle EI, et al. GO::TermFinder--open source software for accessing Gene Ontology information and finding significantly enriched Gene Ontology terms associated with a list of genes. Bioinformatics 20, 3710–3715 (2004).

